# Modulation of the functional interfaces between retroviral intasomes and the human nucleosome

**DOI:** 10.1101/2022.09.29.510092

**Authors:** E. Mauro, D. Lapaillerie, C. Tumiotto, C. Charlier, F. Martins, S. F. Sousa, M. Métifiot, P Weigel, K. Yamatsugu, M. Kanai, H. Munier-Lehmann, C. Richetta, J. Batisse, M. Ruff, O. Delelis, P. Lesbats, V. Parissi

**Author notes:** Viral DNA Integration and Chromatin Dynamics Network (DyNAVir).

## Abstract

Retroviral integration into cell chromatin requires the formation of integrase-viral DNA complexes, called intasomes, and their interaction with the target DNA wrapped around nucleosomes. To further study this mechanism we developed an alphaLISA approach using the prototype foamy virus (PFV) intasome and human nucleosome. This system allowed us to monitor the association between both partners and investigate the protein/protein and protein/DNA interactions engaged in the association with chromatin. Using this approach, we next screened the chemical OncoSET library and selected small molecules that could modulate the intasome/nucleosome complex. Molecules were selected as acting either on the DNA topology within the nucleosome or on the integrase/histone tail interactions. Within these compounds, doxorubicin and histone binders calixarenes were characterized using biochemical, structural and cellular approaches. These drugs were shown to inhibit PFV and HIV-1 integration *in vitro* as well as HIV-1 infection in primary PBMCs cells. Our work provides new information about intasome-nucleosome interaction determinants and paves the way for further unedited antiviral strategies that target the final step of intasome/chromatin anchoring.

## INTRODUCTION

Retroviruses must integrate their viral DNA (vDNA) into the host cell genome to achieve productive infection. This reaction is catalyzed by the intasome nucleoprotein complex, which is composed of an oligomer of integrase (IN) engaging the ends of the vDNA. The first intasome that has been structurally characterized is the spumaretroviral PFV intasome, using X-ray crystallography, revealing a complex composed of four IN protomers (1). It has been generally believed that this tetramer is also sufficient to catalyze the integration of other retroviruses. However, elucidation of intasome structures from other retroviral genera revealed a different organization of these complexes: intasomes from the alpharetrovirus RSV and the betaretrovirus MMTV are composed of an oligomer of 8 INs (2), while the lentivirus MVV and HIV-1 may contain 12-16 INs (3, 4). More recently, another lentiviral intasome, SIV, was revealed to be composed of 12 INs (5). Despite the apparent heterogeneity of these complexes, they share a common core structure called the conserved intasome core, which exhibits a similar organization to the tetrameric PFV intasome. However, these differences in the global quaternary structure also suggest possible differences in their interfaces with cellular partners and nucleosomal substrates that may account for previously observed differences in their sensitivity toward chromatin structure (6–8) and, thus, at least partially, in their preference for host insertion sites. This is further supported by a recent report showing distinct *in vitro* interactions between chromosomes and either the PFV or the HIV-1 intasome (8).

These observations may be in agreement with the distinct cellular insertion sites found for the different retroviruses. Indeed, retroviruses do not integrate randomly into the host cell genome. Numerous cellular cofactors and viral proteins have been shown to be involved in tethering the intasome to specific regions of chromatin, depending on the retrovirus. However, the final target of the integration remains the nucleosome. Nucleosomes are composed of cellular DNA (147 bp) wrapped around an octamer of histone proteins (H2A, H2B, H3 and H4, with 2 copies of each of the histones). To date, structural details of the intasome-nucleosome complex are available only for PFV (9, 10). These structures highlighted a specific positioning of the intasome on the nucleosome, involving several interfaces with three IN subunits, both gyres of the nucleosomal DNA, the core of H2B and the tail of H2A. Similar requirements for direct contact with histone tails have been reported for lentiviral models such as HIV-1 (11, 12). Even if the intasome-nucleosome complex displays multiple interfaces, point mutations in the IN targeting each of these interfaces could drastically reduce both the interaction with the nucleosome and the chromosomes as well as the cellular integration efficiency (9, 11)(13), pointing to them as possible candidates for therapeutic agents or molecular chemical tools.

Thus, the multiple intasome/nucleosome interfaces determine the formation of the actives complexes. A deeper analysis of these interfaces is needed for a better understanding of the dynamic assembly of these supramolecular complexes. With this purpose, we set up an alphaLISA approach to monitor all of the protein/protein and protein/DNA interactions engaged within the intasome/nucleosome complex. A pharmacology approach was then used to select small molecules targeting the specific PFV intasome/nucleosome complex that were able to scan the functional interfaces and modulate its assembly. Several drugs have been identified based on their capability to dissociate the complex by acting either on IN/histone or histone/DNA interactions or on DNA topology. Their mechanism of action was further characterized, leading us to identify new molecular modulators of the intasome/nucleosome functional interfaces and potential new lead compounds for the development of therapeutic agents affecting intasome/nucleosome complex stability.

## RESULTS

### Monitoring the functional PFV intasome/nucleosome complexes using alphaLISA technology

To set up an alphaLISA approach to monitor the complex formed between the PFV intasome and the human mononucleosome (MN), we first assembled both complexes fused to suitable tags. The PFV intasome was assembled following the reported procedure (14) using its cognate vDNA fused to a DIG tag at the 5’ end of the viral ODN mimicking the final ends of the PFV LTR U5 sequence. As shown in **Figure 1A**, the elution profile of the intasome was consistent with published works. The MN was assembled using purified human octamers and a biotinylated 601 Widom sequence (15) following typical salt dialysis procedures, as previously used in the laboratory (8). Each MN assembly was checked using native 8% polyacrylamide gel as reported in **Figure 1B**. Stable native biotinylated MNs obtained with only poorly detectable amounts of free DNA were used for functional and interaction assays. The functionality of the intasome/nucleosome complexes was controlled by a typical concerted integration assay and is shown in **Figure 1C**. A major single integration product resulting from a unique docking position of the intasome onto the MN could be detected, leading to the formation of large (L) and small (S) bands resulting from the previously observed PFV insertion site (9). To better monitor the integration products and fully validate the functionality of the purified intasome, we also assembled an intasome with FITC-coupled viral DNA. No change in the assembly profile was observed when PFV FITC-IN was used (data not shown). As reported in **Figure 1D**, the FITC intasome was also found to be active on MNs, generating the expected integration products and confirming the functional assembly of the complex. An integration assay performed on naked 601 DNA also showed that MN was a preferential substrate and that integration sites detected on MN were specific to this structure. All these data fully confirmed that functional intasomes and MN nucleocomplexes could be assembled using differently tagged partners, allowing their use in alphaLISA and integration assays.

**Figure 1.**
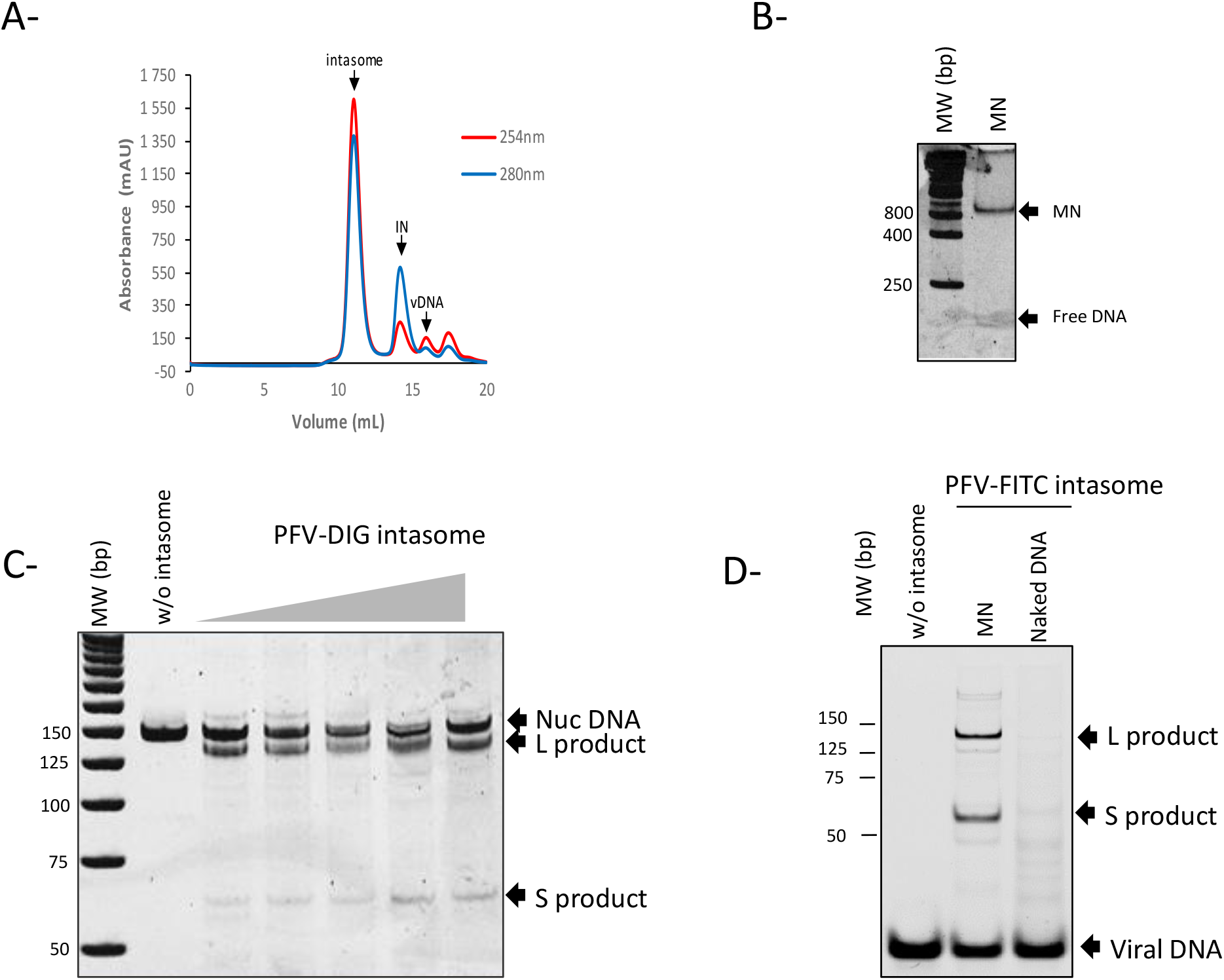
Assembly of functional PFV intasome and human nucleosome. The PFV intasome was assembled as described in Materials and Methods section and subsequently purified by size exclusion chromatography (**A**). Protein and DNA elution was monitored by 280 nm and 254 nm absorbance respectively. MN was assembled by salt dialysis protocol using 5’ biotinylated 601 Widom sequence and human octamers. The structure of the nucleosome was checked by native 5 % polyacrylamide gel (**B**). The functionality of the intasome complex was checked by *in vitro* concerted integration assay performed on the assembled MN using 0-710 nM of intasome and 100ng of MN for 30mn. Integration was monitored either by 1 % agarose gel stained with SybrSafe (**C**). Single site Integration results in the formation of long (L) and short (S) products. Integration catalyzed by FITC-coupled intasome is monitored directly onto gel (**D**).

AlphaLISA technology was employed as shown in **Figure 2A** using anti-DIG acceptor and streptavidin donor beads. The cross titration experiments shown in **Figure 2B** indicate that a strong intasome/nucleosome interaction signal could be detected (>10^6^ counts), even at low concentrations of each partner (6-100 nM). Additionally, titration experiments reported in **SI1** showed that a robust AS signal >10^6^ counts could still be obtained with 1 nM of each intasome and MN partner. These optimal conditions were thus used in the following parts of the work. Control experiments, shown in **Figure 2A**, performed with similar amounts of beads, MN or intasome alone confirmed the specificity of the interaction signal observed with both partners (**Figure 2C**). As reported in **Figure 2D**, the addition of free unlabeled intasome led to an inhibition of the DIG-intasome/Biotin-MN interaction, showing a typical sigmoid competition curve. Fitting of the curve allowed us to calculate the apparent IC_50_ of 27 nM for the complex formed under these conditions. These competitive displacement measurements confirm the specificity of this alphaLiSA binding assay and validate its use to identify compounds that modulate intasome/MN interactions. Since the intasome/MN interactions involved multiple interfaces, including protein/protein and protein/DNA interactions, we compared the alphaLISA signal obtained with the intasome and either the MN or the corresponding 147 bp 601 DNA. A similar interaction signal was detected in the presence of MN and DNA (**SI2**) when using 100 mM NaCl concentrations. However, the sensitivity of the interactions to increasing salt concentrations was found to be different. Indeed, the interaction signal between the intasome and DNA decreased more rapidly than that between the intasome and MN. While a significant interaction signal was still detected at 200 mM NaCl in the presence of MN, the AS signal completely dropped to the zero baseline in the presence of DNA. This confirmed that at concentrations below 200 mM, protein/protein interactions participate in the nucleocomplex interactions in addition to protein/DNA interactions.

**Figure 2.**
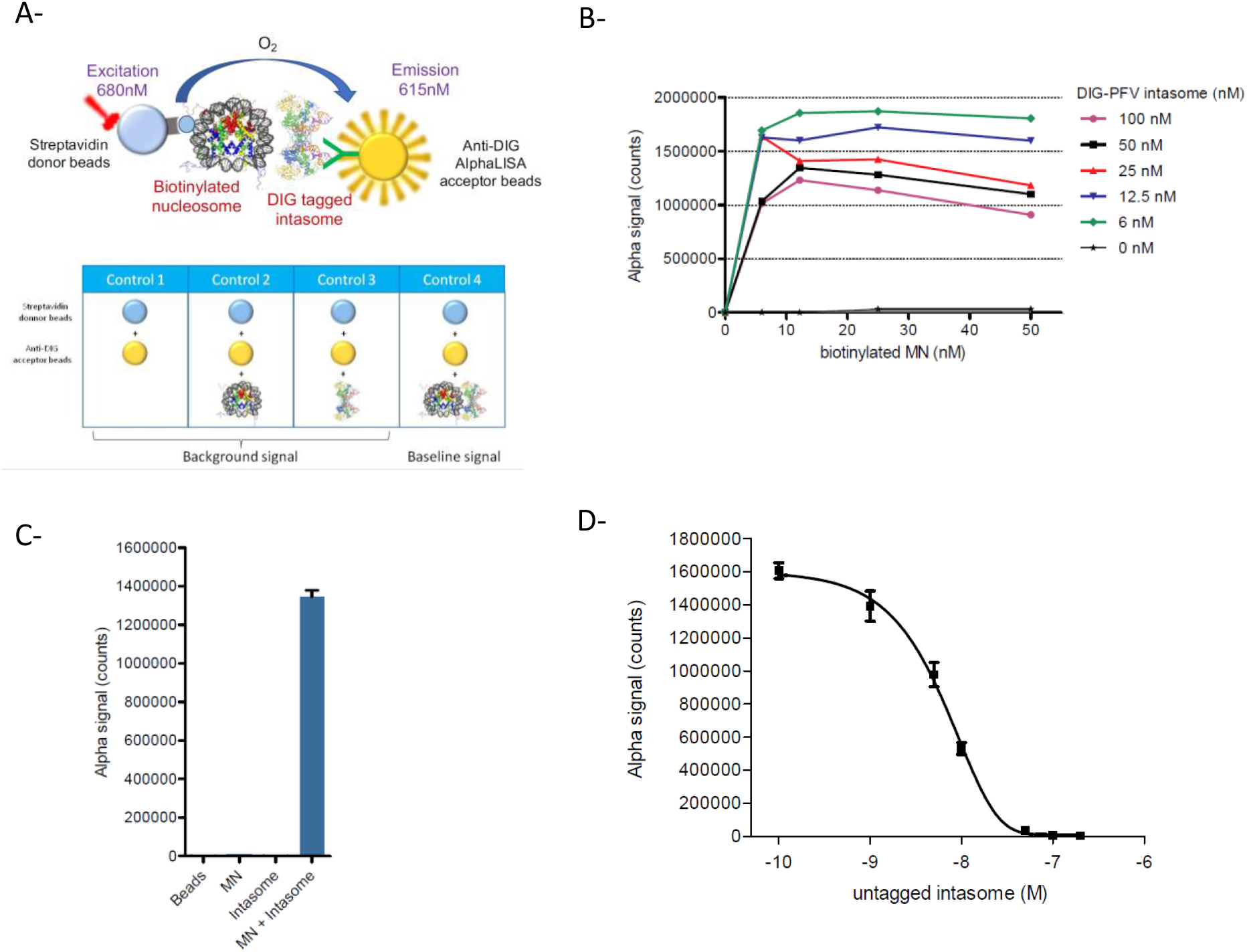
Setup of the intasome/nucleosome AlphaLISA assay. The principle of the AlphaLISA approaches using human biotinylated MN and DIG-tagged PFV intasome as well as the control conditions are schematized in (**A**). The AlphaLISA interaction signal (AU) was monitored using increasing concentrations of each partners and data are reported in (**B**) as a representative experiment. Comparison of the interaction signal obtained with both partners (1 nM), each partner or beads alone is reported in (**C**) as mean of three independent experiments ±SD. Competition experiment performed with increasing concentration of untagged intasome is reported in (**D**) as mean of three independent experiments ±SD.

Taken together, these data indicate that the functional intasome/MN complex can be monitored using the alphaLISA approach, leading to a robust interaction signal even when using low amounts of each partner. This allowed us to further use this system for better characterization of the complex and validated the feasibility of drug screening assays. We thus used this approach to select compounds that may modulate the intasome/nucleosome interaction.

### Selection of drugs affecting the PFV intasome/nucleosome association

The intasome/nucleosome has been shown to involve interactions between the retroviral IN protein and both DNA and histones (9, 11, 12). Thus, compounds expected to dissociate the intasome/MN complex should target these two types of interactions. Since most of the drug assays require the use of DMSO, we first tested the robustness of our system toward concentrations of this solvent. As shown in **SI3**, a decrease in the AS signal was observed only above 2.5% DMSO, and the signal was decreased by 2 at 10% and by 6 at 20%. We concluded that the limit concentration of DMSO usable in this assay was 2.5%. Then, the NIH OncoSET library was screened since it contains up to 133 FDA-approved compounds targeting chromatin-associated proteins and DNA topology that constitute determinants of the intasome/nucleosome complex. Screening was initially performed using 25 μM of each drug in 384-well plates and previously optimized interaction conditions with 1 nM of each partner as described in the Materials and Methods section. Negative controls (beads, MN and intasome alone) were systematically added to the screen. As reported in **Figure 3A**, among the 133 drugs, seven compounds were selected as inducing a decrease of at least 50% in the AS signal. Each selected drug was retested at 10 μM for its effect on the intasome/nucleosome interaction on the AS signal, and as reported in **Figure 3B**, the results confirmed the strong inhibitory effect observed for mitoxanthrone, idarubicin, daunorubicin, doxorubicin and pirarubicin, while tamoxifen showed a 50-60% decrease in the AS signal, and osimertinib showed a 30-40% decrease in the AS signal. As shown in **Figure 3C**, dose‒response sigmoid curves were observed for all drugs except for tamoxifen, which showed only a slight inhibition effect, and osimertinib, which showed no inhibition in the concentration range used and was thus excluded from further analysis. The IC_50_ values of the selected drugs reported in **Figure 3D** were between 0.383 and 37.059 μM. All these compounds were further selected and tested in *in vitro* integration assays.

**Figure 3.**
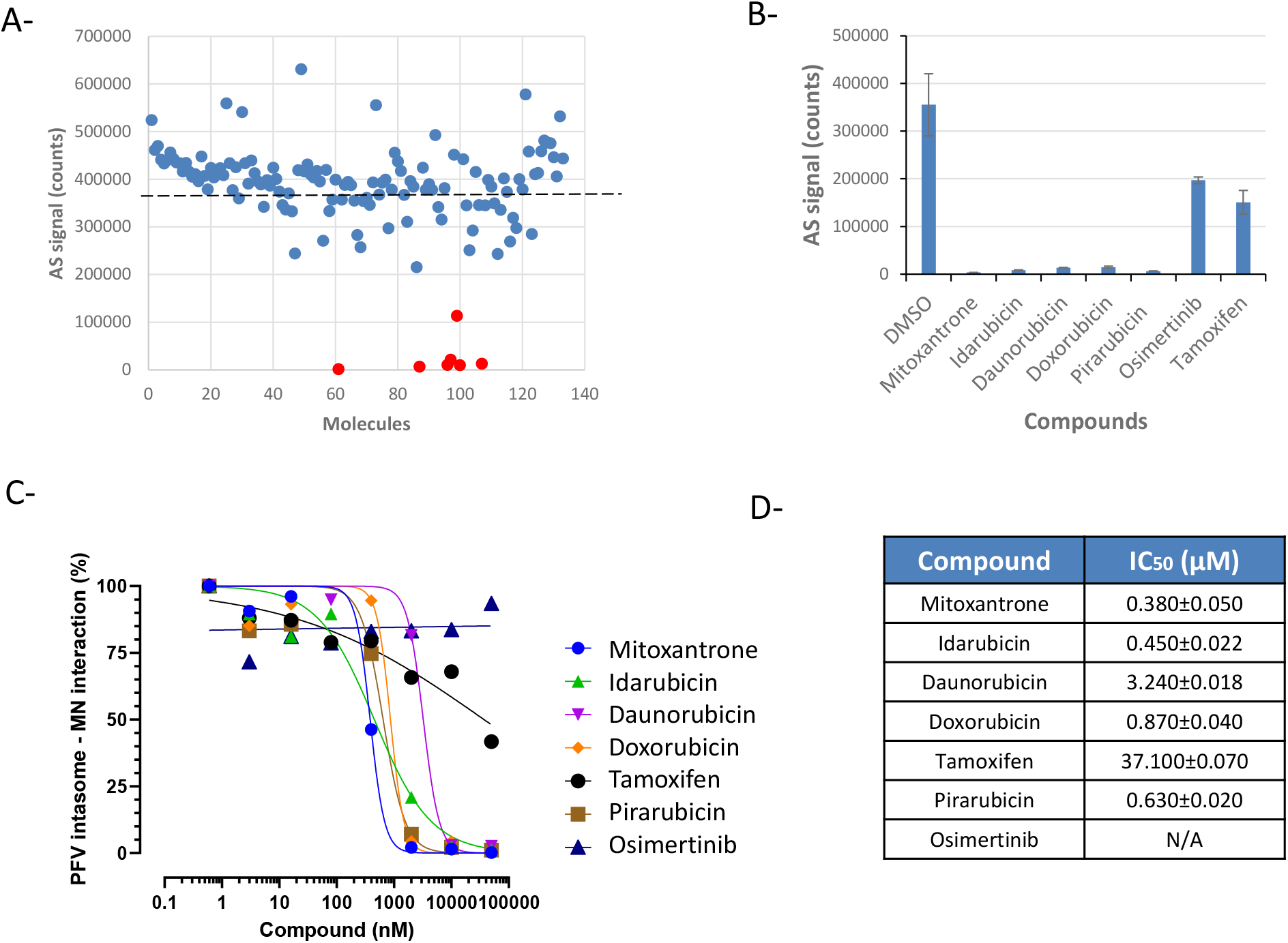
Selection of drugs preventing the intasome/nucleosome association from the ONCOSET NIH library. The 133 compounds of the OncoSET library have been tested at 25μM on the PFV intasome/nucleosome AlphaLisa interaction (**A**). The selected compounds indicated as red bullets have been re-tested at 10 μM (**B**) and the validated drugs have been further tested using increasing concentrations (**C**). The calculated IC_50_ are reported in (**D**). Data are reported as mean of two to three independent experiments ±SD.

### Selection of compounds specifically inhibiting *in vitro* PFV nucleosomal integration

To determine whether the inhibition of the intasome/nucleosome interaction in the alphaLISA assay could be associated with an inhibition of the functional integration reaction, the selected drugs were tested in a typical *in vitro* concerted integration assay. As shown in **Figure 4**, idarubicin, daunorubicin, doxorubicin and pidorubicin, which are the most efficient drugs for inhibiting the AS interaction signal, were also able to inhibit PFV integration into MNs. Tamoxifen was inefficient in inhibiting integration, which is in agreement with its poor effect on the intasome/nucleosome interaction detected in AS. Strikingly, mitoxanthrone did not show any inhibitory effect on integration despite its negative effect on the AS signal, suggesting that it may be a false-positive, as confirmed by counter assay (see **SI4**). Control experiments performed with additional drugs from the library but not selected by AS, such as the DNA intercalating drug thalidomide and the topoisomerase I inhibitor topotecan, did not show any effect, confirming the specific activity of the selected drugs. The intercalating Cis-platin agent was also used as a positive control of inhibition due to its strong DNA intercalating potency and showed only a slight inhibition of integration into nucleosome. Quantitative and comparative integration assays performed using immobilized nucleosomes allowed us to better determine the inhibitory potency of the drugs (**Figure 4B**). In this assay, the selected drugs showed IC_50_ values of MN integration between 0.1 and 1 μM, which align well with their IC_50_ values determined in the AS assay.

**Figure 4.**
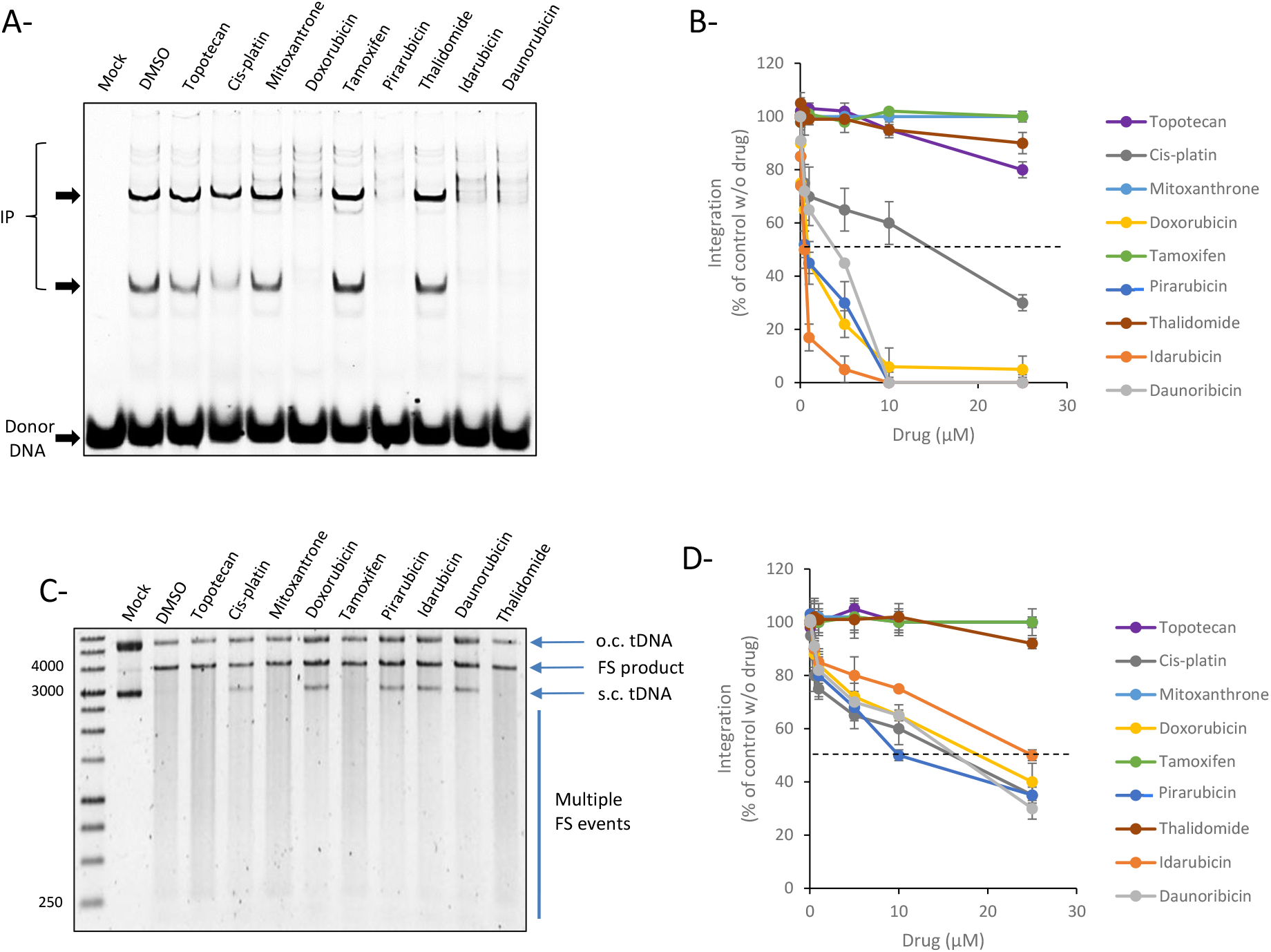
Effect of the selected compounds on *in vitro* PFV integration. A.B. Nuc. C.D. Naked DNA. The compounds selected in **Figure 3** have been tested in typical concerted integration onto mononucleosome using 60nM of FITC PFV intasome and 100ng of MN (**A-B**) or naked DNA plasmid (**C-D**) and increasing concentrations of drugs (0-25 μM). (**A**) and (**C**) show representative experiments performed at 10 μM of drugs and (**B**) and (**D**) show the means from 3 independent experiments ±SD. Dotted line indicates the 50 % of integration.

Integration assays performed on naked DNA plasmids, as reported in **Figure 4C**, showed that most of the drugs exhibited moderate inhibition effects, and the mitoxanthrone and tamoxifen compounds remained inefficient in this assay. Integration into naked biotin-601 immobilized DNA confirmed that the inhibitory effect of the selected drugs was less important on naked DNA than on MN. Indeed, comparison of the effect of the drugs on nucleosomal and naked 601 DNA (**Figure 4B** and **4D**) showed that while the IC_50_ values of the drugs on MN was between 0.1 and 1 μM, the IC_50_ values measured on naked DNA were all above 5-10 μM, suggesting a more specific effect of the drugs on nucleosomal DNA. The more pronounced effect of the drugs observed on the nucleosome led us to further investigate their possible action mechanism related to either the specific nucleosomal DNA structure and/or the histone octamer assembly.

### Effect of the chemical modulation of nucleosomal DNA topology and nucleosome stability by doxorubicin derivates on *in vitro* PFV integration

Based on the previous results, we assume that the drugs affected the target DNA with a preference for the nucleosomal structure. All the best selected compounds were anthracycline enantiomers of doxorubicin known as DNA intercalating drugs (see their chemical structures in **SI5**). Intriguingly, no other DNA intercalating drugs included in the library, such as cisplatin, thalidomide, actinomycin D and bleomycin, were selected, suggesting a specific inhibition mode of the selected doxorubicin derivates. The reported mechanism of action for anthracycline derivates, especially doxorubicin, is binding to DNA intercalated with base pairs, leading to specific binding of the molecules to guanine (as reported extensively in literature). Furthermore, the amino sugar group present in these compounds has also been reported to compete for space with the H4-arginine residues in the nucleosome that may lead to its dissociation (16). Based on their stronger inhibitory effect on MN integration and their DNA intercalating properties, we speculated that the integration inhibition may be due to possible binding to the nucleosomal DNA inducing destabilization of the MN structure. The possible destabilization effect of doxorubicin and its derivative on the MN structure has been further investigated using a previously described histone eviction assay (16). As shown in **Figure 5A**, idarubicin and daunorubicin both induced a strong displacement of histones from the MN, leading to full dissociation of the MN. Doxorubicin and pirarubicin also induced a full shift of the nucleosomal DNA in addition to a strong MN dissociation. To better understand the link between the histone eviction property of the doxorubicin derivate compounds and their integration inhibition property, we took advantage of the previous description of doxorubicinone as an aglycan form of doxorubicin that fails to evict histones from bound nucleosome (16). A histone eviction assay performed with doxorubicin and its aglycan form confirmed that doxorubicinone could not dissociate the MN, in contrast to doxorubicin (**Figure 5B**). As reported in **Figure 5C** and quantified in **5D**, doxorubicinone was also inefficient in inhibiting the *in vitro* integration catalyzed by the PFV intasome onto MNs. The correlation observed between the MN dissociation capability of these drugs and the integration inhibition strongly suggested that their inhibition mechanism was related to the doxorubicin-induced change in nucleosome structure leading to the competition of the amino sugar group of doxorubicin for space with the H4 arginine residue in the DNA minor groove, as previously predicted (16).

**Figure 5.**
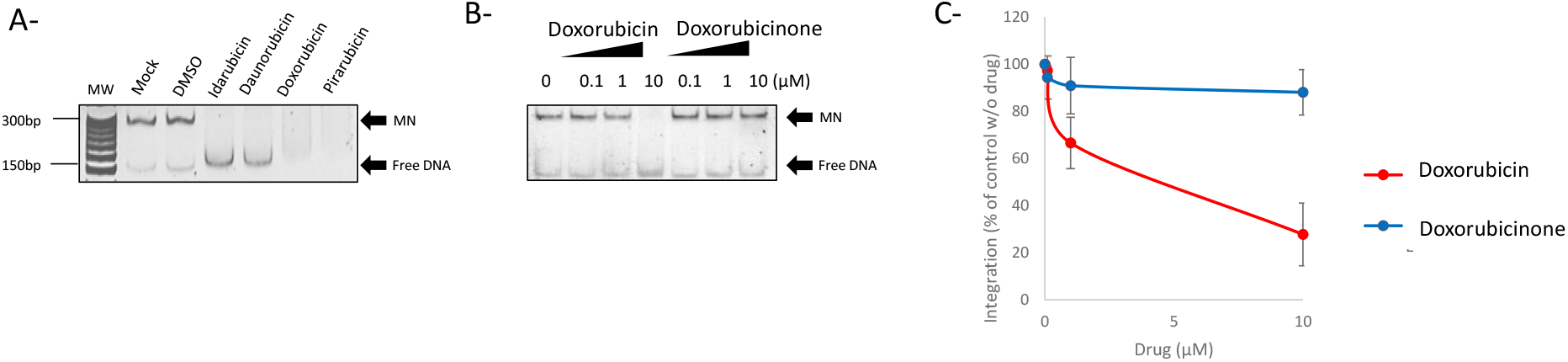
Action mechanism of doxorubicin derivates on nucleosomal integration. The doxorubicin derivates were tested in an histone eviction assay (**A**) using an effective concentration of 10 μM. Doxorubicin and the it’s a-glycan form Doxorubicinone have been compared in histone eviction assay using increasing concentrations of drugs (**B**). The native MN and dissociated naked DNA positions in a SybrSafe stained native 8 % polyacrylamide gel are reported. Doxorubicin and doxorubicinone have been then tested on PFV concerted integration onto nucleosome using increasing concentration of drugs as performed in **Figure 4**. Data are reported as quantification of 3 independent experiments reported as mean ±SD in **D**).

### Computational analysis of the effect of doxorubicin on intasome/nucleosome stability

To better determine the mechanism of doxorubicin inhibition, its interaction with the nucleosome was first modeled using a 50 ns MD simulation as reported in the Materials and Methods section. **Figure 6A** represents the dominant conformation adopted by the doxorubicin molecules in the last 50 ns of the MD simulation. The simulation shows that the doxorubicin molecules interact with the nucleosome mainly at the DNA chains. Interestingly, while the initial MD conformation generated with packmol places the doxorubicin molecules randomly and uniformly distributed along the DNA surface, after the initial 20-50 ns of MD simulation, the doxorubicin molecules tend to pack together and crowd, intercalating between the DNA chains. As evidenced in **Figure 6A-B**, these interactions typically involve two or more doxorubicin molecules, and once formed, they are very stable. **SI6** presents the RMSD of each of the doxorubicin molecules as calculated along the 300 ns of MD simulation. The results show that the conformations adopted by each doxorubicin molecule are quite stable once formed. **Figure 6C-E** shows one of the typical stable interaction modes observed from the MD simulation for the DNA-doxorubicin interaction. It involves an intercalation associated with two parallel doxorubicin molecules placed between the two DNA chains. As illustrated also in **Figure 6A-B**, once formed, this binding mode is very stable and is maintained throughout the remainder of the simulation.

**Figure 6.**
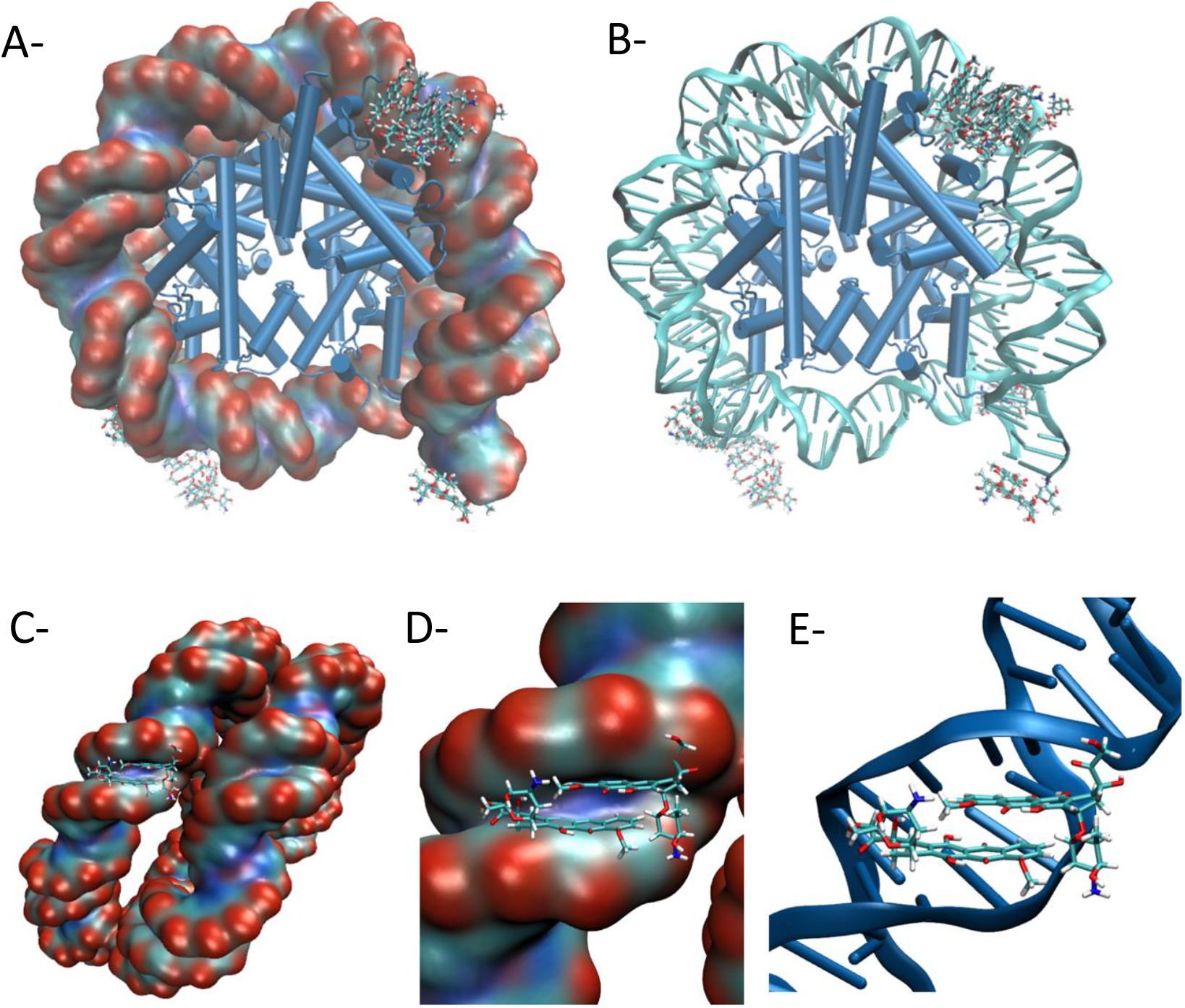
Modelization of the doxorubicin association with the nucleosome. The figure shows the structure of the nucleosome illustrating the overall distribution of the doxorubicin molecules after 300 ns of MD simulation. (**A**) Representation in surface of DNA; (**B**) Representation in cartoon of DNA. In both representations the histones are represented in cartoon and the doxorubicin molecules are represented in licorice. Water molecules were omitted for clarity. The representation of one of the most common doxorubicin binding modes observed through the 300 ns of MD simulation, with two doxorubicin molecules adopting a planar conformation and intercalating between the two DNA helices is reported in (**C**) as a surface-view of DNA, in (**D**) as the detail of the binding surface and (**E**) as a cartoon representation of DNA. In all representations the histones and water molecules are omitted for clarity and the doxorubicin molecules are represented in licorice.

The MM-GBSA calculation was then performed to obtain further information about the intasome/nucleosome dissociation effect of the selected doxorubicin drug. For this purpose, histone-DNA binding free energy was estimated using MM-GBSA, and analyses of the RMSD and RMSF of the 300 ns MD simulations on the systems were performed with and without doxorubicin.

We first analyzed the impact of doxorubicin binding on nucleosome energetic stability. The protein‒DNA binding free energy ΔG_bind_ was estimated by MM-GBSA calculation. Values of the individual components are presented in **Table 1**, including the gas-phase electrostatic (Δ*E*_ELE_) and van der Waals (Δ*E*_VDW_) interaction energies, the polar solvation energy (ΔG_GB_) calculated by using the generalized-born (GB) model, and the nonpolar solvation energy (Δ*G*_Surf_). ΔG_Polar_ is written as the sum of the polar energy terms, which include Δ*E*_ELE_ and ΔG_GB_, while ΔG_Non-Polar_ represents the sum of the nonpolar-based energy terms, which include Δ*E*_VDW_ and Δ*G*_Surf_. As reported in **Table 1**, the protein‒DNA binding free energy of -970.9 kcal/mol in the absence of doxorubicin dropped to a less stable binding free energy of -937.8 kcal/mol upon the addition of doxorubicin, leading to a predicted decrease in stability of 33.1 kcal/mol (3.4%). Analysis of the different individual contributions to the binding free energy, as calculated using the MM-GBSA method, showed that this difference arose from a decrease in both the polar (15.5 kcal/mol) and nonpolar components (17.5 kcal/mol), i.e., it has a 47% polar/53% nonpolar contribution. More detailed analysis of ΔΔ*E*_ELE_, ΔΔ*E*_VDW_, ΔΔG_GB_, and ΔΔ*G*_Surf_ upon doxorubicin binding unambiguously showed that the doxorubicin interaction decreased the direct electrostatic and van der Waals interactions between protein and DNA. Consequently, on the basis of these calculations, doxorubicin binding to MN should lead to a less stable structure.

**Table 1.** Nucleosome Histone-DNA binding free energies estimated by MM-GBSA in the absence and in the presence of doxorubicin. All values in kcal/mol.

| Model | $\Delta G_{\text{bind}}$<br>(kcal/mol) | $\Delta E_{\text{ELE}}$<br>(kcal/mol) | $\Delta E_{\text{VDW}}$<br>(kcal/mol) | $\Delta G_{\text{GB}}$<br>(kcal/mol) | $\Delta G_{\text{Surf}}$<br>(kcal/mol) | $\Delta G_{\text{Polar}}$<br>(kcal/mol) | $\Delta G_{\text{Non-Polar}}$<br>(kcal/mol) |
| --- | --- | --- | --- | --- | --- | --- | --- |
| Without Doxorubicin | -970.9 ± 1.0 | -163,522.1 ± 16.7 | -764.3 ± 0.5 | -163,422.2 ± 16.4 | -106.6 ± 0.1 | -99.9 | -870.9 |
| With Doxorubicin | -937.8 ± 0.9 | -163,161.3 ± 17.6 | -748.6 ± 0.6 | -163,076.9 ± 17.3 | -104.8 ± 0.1 | -84.4 | -853.4 |
| Variation | + 33.1 | +360.8 | +15.7 | -345.3 | +1.8 | +15.5 | +17.5 |

We then analyzed the impact of doxorubicin binding on nucleosome structural stability. **Table 2** presents the average RMSD of the last 200 ns of the MD simulations performed in the absence and in the presence of doxorubicin in comparison with the nucleosome X-ray structure. The results show that the average RMSD of the backbone atoms in the nucleosome is larger in the simulation performed in the presence of doxorubicin (2.77 Å) than in the simulation performed in the absence of doxorubicin (2.67 Å), illustrating that the presence of doxorubicin induces a structural change. Analyzing the behavior of the protein and DNA components of the nucleosome, it can be observed that this effect is stronger in DNA than in histones. In fact, DNA exhibits a larger RMSD change with doxorubicin addition (from 3.33 Å to 3.58 Å) than the histones do (from 1.43 to 1.48 Å). This was confirmed when regarding the RMSD change over time for both simulations, with and without doxorubicin (**SI6**) showing the same average tendency described for **Table 1**: the RMSD is larger for the simulation in the presence of doxorubicin than it is for the simulations in the absence of doxorubicin; and the RMSD change with doxorubicin addition is larger for DNA than it is for the histones. **SI6** also demonstrates that both simulations are well equilibrated after the initial 50 ns.

**Table 2.** Average RMSd of the last 200 ns of the MD simulations in comparison with the initial nucleosome structure, in the absence and in the presence of doxorubicin.

| Model | Average RMSd Nucleosome<br>(Å) | Average RMSd Histones<br>(Å) | Average RMSd DNA<br>(Å) |
| --- | --- | --- | --- |
| Without Doxorubicin | 2.67 ± 0.13 | 1.43 ± 0.07 | 3.33 ± 0.18 |
| With Doxorubicin | 2.77 ± 0.14 | 1.48 ± 0.08 | 3.58 ± 0.22 |

The impact of doxorubicin binding on nucleosome structural flexibility was then investigated. **Table 3** presents the average root-mean-square fluctuation (RMSF) of the backbone atoms in the nucleosome, illustrating the positional variability/flexibility of the atoms in the nucleosome in the presence and absence of doxorubicin. The results show that the addition of doxorubicin alters the flexibility of the nucleosome. On average, doxorubicin induces a slight increase in the flexibility of the nucleosome residues (1.12 vs. 1.09 Å). This effect is more significant for DNA (1.62 Å with doxorubicin vs. 1.51 Å without doxorubicin) than it is for histones. **SI7** illustrates the RMSF for the different amino acid and nucleotide positions along the histone and DNA chains in the nucleosome in the absence and presence of doxorubicin. The results show a similar profile in terms of relative flexibility for both models. In general, the most flexible amino acid positions and DNA positions are the same, independent of the presence of doxorubicin. However, the simulation in the presence of doxorubicin shows an increase in flexibility, which is particularly noticeable for the DNA portion of the nucleosome. These changes are not evenly distributed but are stronger along specific positions in the nucleosome. **SI8** represents the individual change in RMSF with the addition of doxorubicin. The results confirm the existence of general light flexibility variations along the histone residues, with more significant increases along the amino acid positions/histones 127A-135A, 29D-47D, 134E-25F and 30H-32H. Decreases in flexibility were also observed along positions 19B-22B, 91F-96F and 97G-100G. RMSF changes along the DNA positions upon doxorubicin interaction were much more dramatic and particularly evident among positions -72/-57, -52/-22, and 67/72 in chain I and -72/-63 and +31/+68 in chain J. These regions are more affected by the interaction with doxorubicin molecules.

**Table 3.** Average RMSF of the two nucleosome and of the histone and DNA components in the absence and in the presence of doxorubicin.

| Model | Average RMSF Nucleosome (Å) | Average RMSF Histones (Å) | Average RMSF DNA (Å) |
| --- | --- | --- | --- |
| Without Doxorubicin | 1.09 ± 0.46 | 0.92 ± 0.39 | 1.51 ± 0.35 |
| With Doxorubicin | 1.12 ± 0.54 | 0.93 ± 0.42 | 1.62 ± 0.51 |

### Selection of drugs targeting the protein/histone interaction as inhibitors of PFV integration

In addition to protein/DNA interactions targeted by doxorubicin drugs, protein/histone interactions should also constitute important components of retroviral intasome/nucleosome stability, as previously suggested (9, 11, 12). Tetrasuflonated calix[4]arene molecules have been previously reported to specifically bind to histone tails with submicromolar affinity and strong specificity ((17) and **Table 4**). Among these compounds, CA3 (see the chemical structure in **SI9**) showed the best histone binding affinity and was proposed to be effective for disrupting interactions between histone tails and effector proteins (17). These molecules were thus also included in the intasome/nucleosome AS assay to evaluate their capability to dissociate or block the formation of the PFV intasome/MN complex. As reported in **Figure 7A**, among the calixarene drugs, the CA3 compounds were shown to prevent the formation of the intasome/MN complex in AS, leading to its complete dissociation at concentrations greater than 5 μM. CA2 and CA1 analogs were found to be less efficient in inhibiting the association, which aligns well with their lower histone affinity (see their previously measured Kd for histone tails in **Table 4**).

**Table 4.** Comparison between the histone tails affinity and *in vitro* HIV-1 integration inhibition of CA molecules.

| CA molecule | Kd ($\mu\text{M}$) to histone tails<br>(Kimura et al.,<br>ChemBioChem 2015) | Integration IC<sub>50</sub> |
| --- | --- | --- |
| CA1 | 17.8-20-4 | >1$\mu\text{M}$ |
| CA2 | 1.8 | 0.9 $\mu\text{M}$ |
| CA3 | 0.73-0.51 | 0.1-0.4 $\mu\text{M}$ |

**Figure 7.**
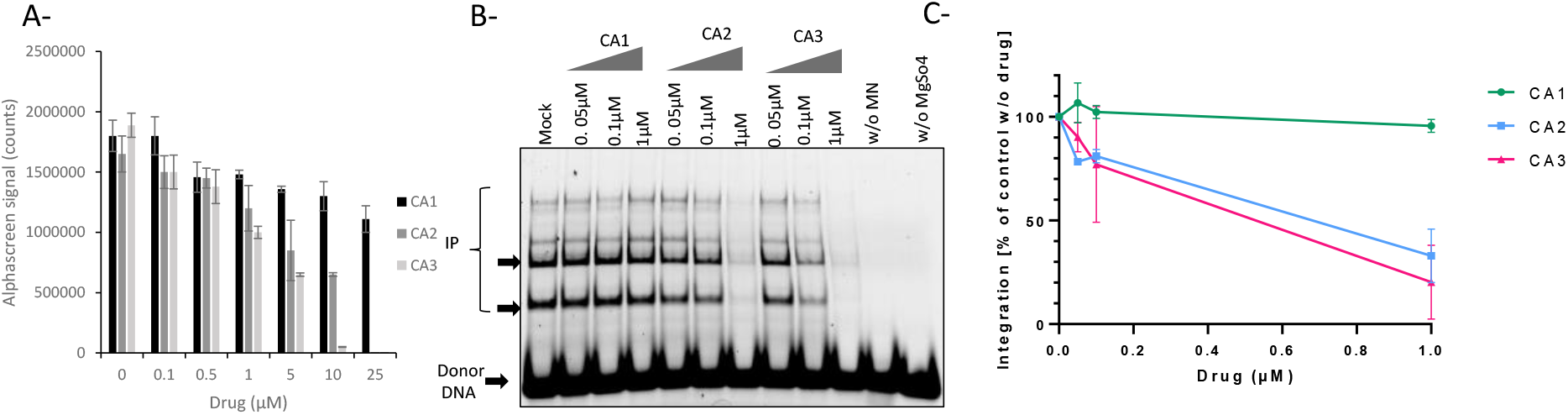
Effect of calixarenes compounds on PFV intasome/nucleosome complex formation and *in vitro* integration. CA1, CA2 and CA3 molecules have been tested on the PFV intasome/nucleosome complex using the AlphaLISA technology as setup in this work (**A**). The drugs were then assayed in a typical concerted integration using FITC-PFV intasome and nucleosome and the integration products were monitored on 8 % native polyacrylamide gel (**B**). The results are reported in (**C**) as the mean of 3 independent integration experiment ±SD.

An *in vitro* concerted integration assay performed on MNs confirmed that CA3 inhibited viral DNA insertion catalyzed by the PFV intasome (**Figure 7B**). Comparison of the inhibitory effect of CA1, CA2 and CA3 on *in vitro* integration catalyzed by the PFV intasome also showed different efficiencies (**Figure 7C)**. Indeed, while CA3 showed a stronger inhibitory effect with IC_50_∼100-400 nM, CA2 showed a lower inhibition capability (IC_50_ ∼900 nM); CA1 was inefficient in our assay. Interestingly, the inhibition efficiency of the drugs closely paralleled their affinity for histone tails, suggesting that the inhibition mechanism was related to these histone components of the nucleosome.

### Effect of the selected compounds on *in vitro* HIV-1 integration

To better address their specificity, the best selected drugs were further tested for HIV-1 integration using an *in vitro* integration assay and performed on MNs with preformed functional vDNA•IN•LEDGF/p75 intasome complexes as previously described (13). As reported in **Figure 8A** and **B**, doxorubicin was also shown to inhibit the *in vitro* integration of HIV-1 with similar efficiency to that shown by the PFV model (IC_50_∼300 nM,). Again, the doxorubicinone derivative was found to be inefficient, strongly suggesting that the mechanism of action of doxorubicin was similar to that elucidated in the PFV model, *i*.*e*., nucleosome dissociation by histone eviction (see **Figure 5-6**). Integration assays performed with the CA3 compound showed that it was also efficient in inhibiting HIV-1 *in vitro* integration (**Figure 8C**). Comparison between the inhibition efficiency of CA1, CA2 and CA3 showed a good correlation between the histone tail binding property of the drugs and their integration inhibition efficiency, with CA3 being the most efficient compound with an IC_50_ of approximately 100 nM (see the determination of the IC_50_ for all molecules in **Figure 8D**). Based on their affinity for histone tails, the inhibition mechanism was assumed to be a competition between the intasome and these tails. This was confirmed by an integration inhibition assay performed on either native MN, tailless MN or naked DNA showing that deleting the tails or using naked DNA increased the IC_50_ of the drug from 0.035-0.092 μM to 0.127-0.913 μM and 0.178-0.810 μM, respectively (**SI10**). These results confirmed that the main mechanism for the integration inhibition mediated by CA3 involves competition with intasome binding to the histone tails.

**Figure 8.**
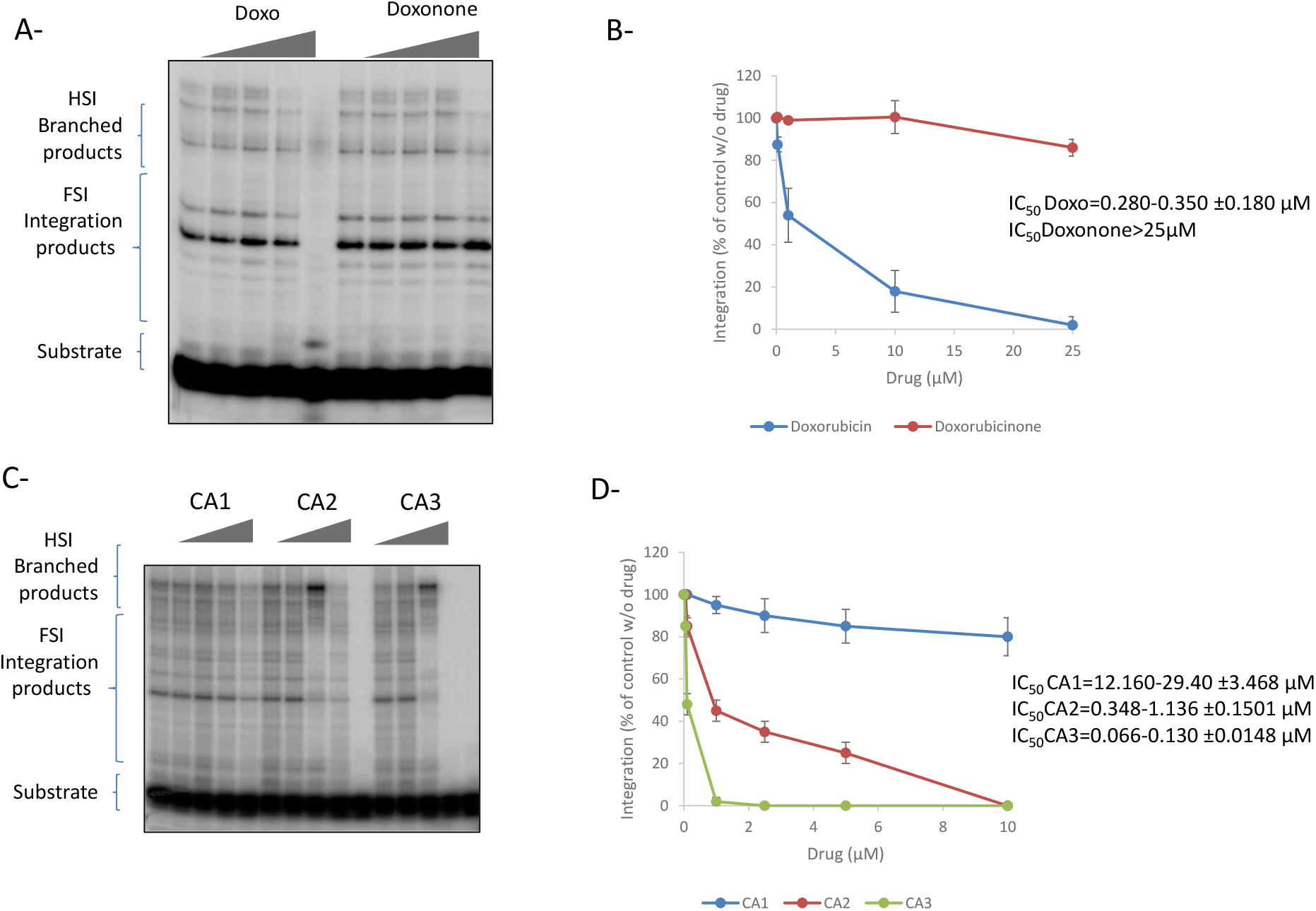
Effect of the selected drugs on *in vitro* HIV-1 integration. Doxorubicin /doxorubicinone (**A, B**) or CA1/2/3 (**C, D**) have been added to a typical *in vitro* concerted integration performed with preformed HIV-1 intasome, radiolabeled viral U5 end fragment and mononucleosome. The integration products were monitored on 6-12 % gradient polyacrylamide gel (left panels) and quantified (right panels). Data are reported as mean of 2-4 independent experiments ±SD and IC_50_ were reported in the figure.

### Effect of the selected compounds on *in cellulo* HIV-1 replication

Next, we further address the efficiency of the drug in a cellular context. We first checked the cytotoxicity of the compounds in different cell lines to evaluate their possible use as antiviral agents. MTT cytotoxicity assays performed with the drugs selected using the AS screen of the OncoSET library, including the doxorubicin derivatives, showed that they were all cytotoxic in typical transformed or cancer cellular models within different concentration ranges (see **SI11**). This was expected since these compounds are all anticancer drugs that act by inducing the death of cancerous cells. In contrast, a poor effect was observed in noncancerous primary PBMCs. In contrast to doxorubicin derivatives, the CA molecules were shown to have little to no cytotoxicity in our assays in all the cell lines, including PMBCs (see **SI11**). Based on these data, both selected drugs were assayed for HIV-1 infection in primary PBMCs.

Infection assays performed with the clinical B subtype HIV-1 strain in PBMCs treated with doxorubicin showed a strong inhibition of replication by the drug, leading to an IC_50_ of 2-20 nM (**Figure 9A**). The CA3 compound was also found to decrease viral replication with an IC_50_ of 1-2 μM (**Figure 9B**). qPCR quantification of the total DNA at 20 h postinfection showed no effect of the Doxotubicin, indicating that the drug does not affect the reverse transcription or the entry step (**Figure 9C**). In contrast, integrated DNA was found to be decreased in doxorubicin-treated cells, while two-LTR circle amounts were increased (**Figure 9C**). These data confirmed that doxorubicin inhibited the postnuclear integration steps. Quantification of the total DNA in infected cells treated with CA3 showed no effect of the drugs on RT at concentrations up to 2 μM, where a severe decrease in replication was observed (**Figure 9D**). However, higher concentrations (20 μM) induced a slight decrease in total DNA synthesis, suggesting that under these concentrations, the drugs might affect reverse transcription or virus entry. The integration inhibition was confirmed by quantification of the integrated and unintegrated circular DNA showing the expected signature of integrated DNA increase accompanied by an increase in two-LTR circles.

**Figure 9.**
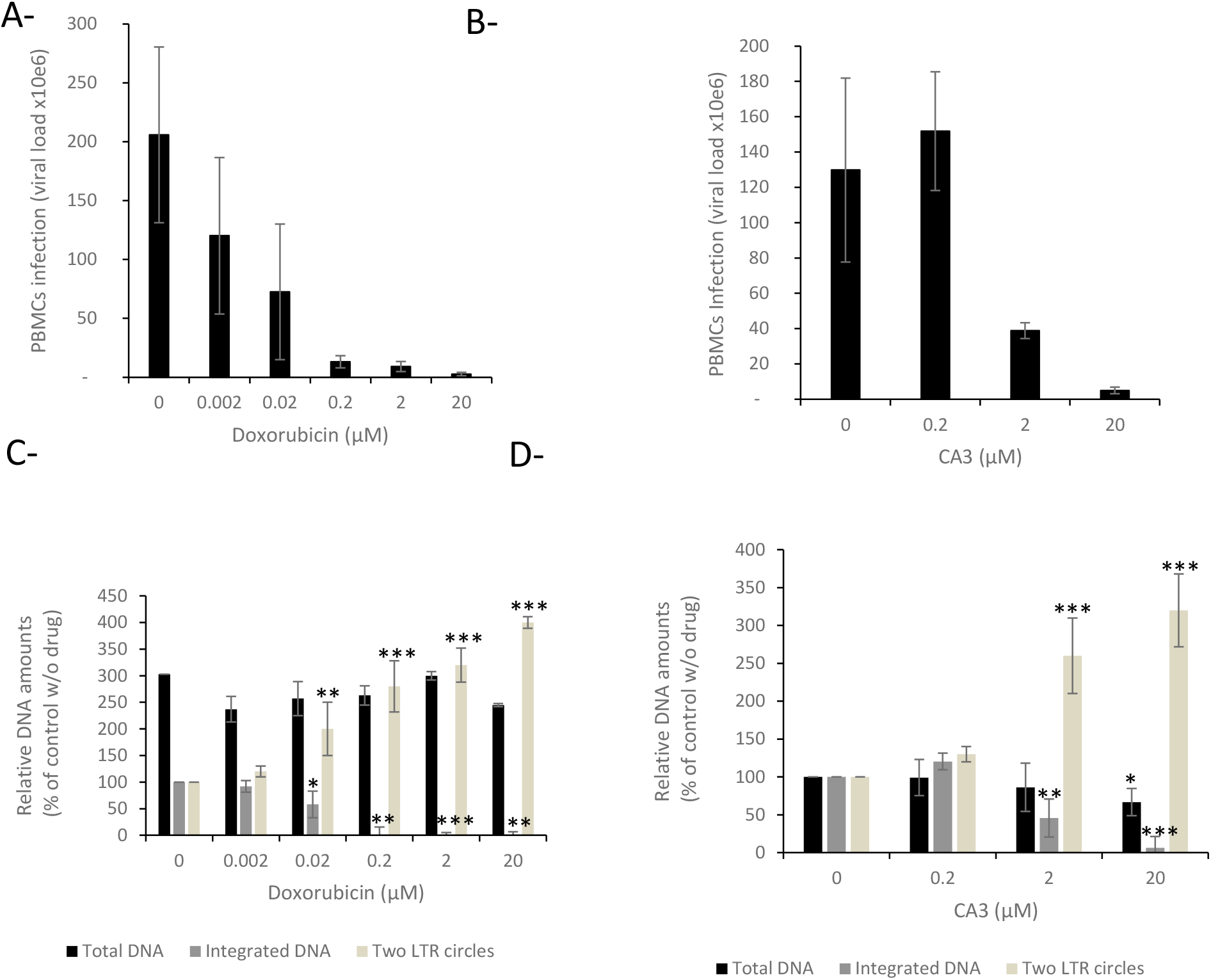
Effect of the calixarenes compounds of HIV-1 replication. PBMCs cells have been infected with transduced with lentiviral vector in the presence of increasing concentration of CA1/2/3 drugs. Infectivity was measured as percentage of GFP positive cells and reported as percentage of control without drug (**A**). Viral DNA populations have been quantified by qPCR (**B-C**). Data are reported as mean of 2-4 independent experiments ±SD.

Taken together, our data suggest that the CA3 histone binder and doxorubicin can prevent integration in infected cells and may serve as tools for further dissecting the retroviral intasome/chromatin interaction and as candidates for further therapeutic developments.

## CONCLUSIONS

The retroviral intasome/nucleosome complex is the functional entity responsible for the stable integration of the viral genome into the host chromatin. Structural data about this complex are available only for the PFV model (9). These data show that the intasome/nucleosome complex involves multiple types of interactions, including protein‒ protein and protein‒DNA interactions within either the nucleosome, the IN-IN oligomer or the intasome-nucleosome complex (9). Chemical modulation of these interfaces is an original way to better understand the role of these interactions in the integration process as well as helping the development of potential molecular tools or therapeutic agents. Both the understanding of the functional architecture of this complex and the search for new inhibitory strategies are required to monitor all these IN/nucleosome interfaces. To this end, we have developed an alphaLISA-derived approach allowing us to recapitulate all the interactions engaged in the retroviral intasome/nucleosome complex, including the IN/target DNA, IN/viral DNA, IN/IN and IN/histone bonds. In addition to providing an experimental model for further depicting the role of these interactions, our approach was used to select new molecules that dissociate the functional complex by targeting each or all of these interfaces.

Since chromatin is a target for anticancer therapy, the OncoSET library was first screened. Among the 133 drugs in the OncoSET database, four were selected as significant inhibitors of both the AS intasome/MN interaction signal and *in vitro* integration. Most of the selected drugs were anthracycline derivates, including doxorubicin, and are known as DNA intercalating agents. Interestingly, inhibition of HIV-1 integrase activity by doxorubicin has been previously reported (18). However, while the inhibition of partial strand-transfer activity onto naked short DNA has been shown with IC_50_ values between 1-10 μM, we reported here a new mechanism of action for this molecule acting specifically on the physiological nucleosomal DNA substrate with IC_50_<1 μM. Interestingly, not all the intercalating agents represented in the OncoSET library, such as actinomycin D, showed AS signal inhibition effects. More strikingly, among the anthracyclines in the OncoSET database, not all were selected. For instance, epirubicin was not selected as an intasome/nucleosome modulator, in contrast to doxorubicin, idarubicin, pidorubicin and daunoribicin, suggesting a specific action mechanism for these anthracycline derivatives, leading us to further investigate the nucleosome dissociation property of doxorubicin as a mechanism for the reported inhibition. The nucleosome dissociation property of this molecule has been previously reported to be due to the competition of the amino sugar group of the drug for space with the H4-arginine residue in the DNA minor groove, since this H4-DNA interaction stabilizes the nucleosome structure (19). Both computational and biochemical analyses of the mechanism of action of the drug confirmed that the main process leading to integration inhibition was the dissociation of the nucleosome structure by the histone eviction induced by the competition for the space with the H4-arginine residues in the nucleosome mediated by the amino sugar group present in these compounds. The lack of effect observed with other strong DNA intercalating agents, such as actinomycin D, also suggested that the dissociation of the nucleosome induced by the binding of drugs to DNA is the prerequisite for efficient integration inhibition of this substrate due to the destabilization of the MN structure by affecting the nucleosomal DNA topology. This was further demonstrated by computational MM-GBSA analyses confirming that the binding of doxorubicin on the nucleosomal DNA induces physical constraints both at the DNA and protein levels, leading to the destabilization of the complex. Taken together, these data suggest a new integration inhibition process specifically targeting the physiological nucleosome substrate by anthracycline derivates. The results also confirm the importance of the fully assembled nucleosome for efficient integration, shedding light on previous observations showing that the remodeling process favors HIV-1 integration (6, 20). Indeed, altogether, these data suggest that enhancement of integration may require the remodeling of neighboring chromatin, allowing the intasome to gain access to a fully native targeted nucleosome or poly-nucleosome at the insertion site. Integration inhibition observed in HIV-1-infected primary cells confirms that these interactions are also important in the physiological complex.

With the same methods of probing the protein/DNA interfaces, we also wanted to evaluate the IN/histone interaction importance by chemical targeting using histone binders as previously reported calixarene molecules (17). The alphaLISA approach led us to show the inhibition of the intasome/nucleosome interaction by the CA3 drug. *In vitro* integration assays further demonstrated that this drug was able to inhibit nucleosomal integration. An assay on other retroviral integration systems, such as HIV-1, confirmed that the selected drugs could also be efficient in inhibiting this integration process *in vitro*, further validating the approach for selecting drugs aimed at targets of potential therapeutic interest. Comparison of the effect of the CA3 drug and chemical analogs with less affinity for histone tails, in addition to assays using tailless nucleosomes, confirmed the histone-dependent inhibition mechanism of the drug. These results validated the chemical dissociation of the IN/histone tail interaction as a new retroviral inhibition mechanism. These results parallel those of previously published data showing that calixarene compounds could inhibit histone binding by histone readers both *in vitro* and *in cellulo* (21, 22). In this line, our work paves the way for the possible development of calixarenes derivates specific for some modifier histone tails possibly recognized by retroviral intasomes (such as the H3K36me3 modification of lentiviral intasomes assumed to be associated with LEDGF/p75, or the histone H4 tail bound by HIV-1 or H4ac associated with BET proteins participating to the chromatin anchoring of gammaretroviral intasomes) (23, 24). Our results also confirm the importance of intasome/histone interactions for efficient HIV-1 integration, as previously suggested (11, 12) and demonstrated for the PFV model (9).

In addition to serving as interesting tools for dissecting the molecular processes involved in the functional anchoring of the intasome to the nucleosome, we wondered whether the selected compounds could also be the foundation of new therapeutic approaches. To this end, we further studied the inhibitory capability of the selected drugs on the HIV-1 model of direct therapeutic interest, focusing on doxorubicin and CA3. Both doxorubicin and CA3 compounds showed poor cytotoxicity in primary PBMCs up to 38 μM, which allowed us to investigate their effect on viral infectivity. The data indicated that the two drugs were able to inhibit HIV-1 infectivity with IC_50_ values between 2 nM and 1-2 μM. Quantification of the viral integrated and unintegrated DNA populations confirmed that the drug was able to specifically block the integration step.

These data show that the selected drugs are good candidates for new antiviral lead compounds and provide a new antiviral precept and may pave the way for further developments of such strategies targeting the integration step. Additional work will be required for optimizing the inhibitory property of the drugs as well as improving the cellular tolerance of them. The selected drug may serve for further structure-activity (SAR) studies to both improve its inhibition efficiency and specificity as well as decrease possible cellular toxicity. To this end, our study provided some information for these future improvements. Indeed, epirubicin is a stereoisomer in which the hydroxyl group of doxorubicin is inverted at position 4’ and has a similar mechanism of action as that of doxorubicin, while the compound is inefficient in inhibiting the intasome/nucleosome complex. This may underline the importance of position 4’ in the inhibition mechanism. The inefficiency of valrubicin may also be taken into account for SAR studies. The biochemical data as well as the *in silico* GBSA modelling provided in our work will also serve as a basis for such improvement. The validation of the alphaLISA reported here also opens the way for broader screening using larger libraries that should allow the identification of additional compounds modulating the intasome/nucleosome interfaces important for the integration process. Indeed, no drugs targeting the intasome alone have been selected, probably due to the nature of the selected library encompassing mainly drugs targeting the chromatin components. An additional library may circumvent this issue and lead to the selection of additional inhibition processes.

In addition to identifying compounds interfering with intasome/nucleosome binding, our data confirm that the AS approach reported here constitutes a robust model for selecting drugs able to modulate the intasome/nucleosome interfaces in several retroviral models, including those of therapeutic interest. Both anthracycline derivatives and the histone binder CA3 selected in this system using the PFV model were active in inhibiting *in vitro* HIV-1 integration, confirming that the PFV intasome/nucleosome alphaLISA approach is suitable for selecting potential anti-HIV compounds.

Our results also prove that alphaLISA is a technology also suitable for bimolecular inhibitor screening assays using large complexes such as intasomes (several hundreds of kDa) and nucleosomes (∼200 kDa). This approach has the additional advantage of recapitulating all the interfaces engaged in these large nucleocomplexes, allowing the selection of molecules that may target all these interfaces, in contrast to approaches based on minimal partners. In our study, the AS approach particularly allowed us to identify drugs targeting either the MN tail or the DNA. The developed model and the selected drugs will help in better dissecting the molecular interactions within the intasome/nucleosome functional complex. In particular, this first monitoring of the retroviral/intasome complex paves the way for multiple applications, including the analysis of the influence of cellular partners and the study of additional retroviral intasomes, allowing us to determine the possible specific interfaces previously suggested (7, 8, 25). Our work also provides the technical basis for larger-scale screening of drugs specifically targeting these functional nucleocomplexes or additional nucleosome-partner complexes.

## Materials and methods

### Proteins

Intasomes of spumavirus PFV, betaretrovirus MMTV and lentivirus MVV have been assembled as described in. Briefly, IN and their cognate vDNA were mix and assembled intasomes were purified by size exclusion chromatography. Elution profiles of the different intasomes used in this study are shown in **Figure 1A** with their respective structures depicted in **Figure 1B**. Concordantly with published works, the smallest one (PFV) which is composed of a tetramer of IN, eluted around 11 mL while the octameric MMTV eluted at 10.5 mL and finally the hexadodecameric MVV intasome at 8 mL. The HIV1-IN was expressed in E.coli (Rosetta) and the cells were lysed in buffer containing 50 mM Hepes pH 7.5, 5 mM EDTA, 1 mM DTT, 1 mM PMSF. The lysate was centrifuged and IN extracted from the pellet in buffer containing 1 M NaCl, 50 mM Hepes pH 7.5, 1 mM EDTA, 1 mM DTT, 7 mM CHAPS. The protein was then purified on butyl column equilibrated with 50 mM Hepes pH 7.5, 200 mM NaCl, 1 M ammonium sulfate, 100 mM EDTA, 1 mM DTT, 7 mM CHAPS, 10 % glycerol. The protein was further purified on heparin column equilibrated with 50 mM Hepes pH 7.5, 200 mM NaCl, 100 mM EDTA, 1 mM DTT, 7 mM CHAPS, 10 % glycerol. LEDGF/p75 was expressed in PC2 bacteria and the cells were lysed in lysis buffer containing 20 mM Tris pH 7.5, 1 M NaCl, 1 mM PMSF added lysozyme and protease inhibitors. The protein was purified by nickel-affinity chromatography and the His-tag was removed with 3C protease, 4°C over night. After dilution down to 150 mM NaCl, the protein was further purified on SP column equilibrated with 25 mM Tris pH 7.5, 150 mM NaCl (gradient from150 mM to 1 M NaCl), then DTT was added to 2 mM final and the protein was concentrated for Gel filtration. Gel filtration was performed on a superdex 200 column (GE Healthcare) equilibrated with 25 mM Tris pH 7.5, 500 mM NaCl. Two mM DTT were added to eluted protein that was then concentrated to about 10 mg/ml.

### Drugs

Molecules were screened from the OncoSET library (NIH). The selected compounds were then purchased from various private company. The CA compounds were synthetized as previously described (17).

### Integration assays on nucleosome using the intasomes

Integration assays using purified assembled intasomes were performed as followed: 200 ng of mononucleosome were incubated with 30 nm final concentration purified intasome in 100 mM NaCl, 20 mM BTP pH 7, 12.5 mM MgSO4 in a final volume of 40 μL for PFV intasome, and for HIV intasome in 20 mM HEPES pH 7, 7.5 % DMSO, 8 % PEG, 10 mM MgCl2, 20 μM ZnCl2, 100 mM NaCl, 5 mM DTT final concentration. The mix was then incubated at 37°C for 15mins and 1 hour respectively. Then reaction was stopped by the addition of 5.5 μL of a mix containing 5 % SDS and 0.25 M EDTA and deproteinized with proteinase K (Promega) for 1 hour at 37°C. Nucleic acids were then precipitated with 150 μl of ethanol overnight at −20°C. Samples were then spun at top speed for 1 hour at 4°C, and the pellets were dried and then re-suspended with DNA loading buffer. Integration products were separated on an 8 % native polyacrylamide gel.

### AlphaLISA

AlphaLISA is a technology particularly suitable for bi-molecular inhibitor screening assays using protein-protein interactions with purified recombinant proteins. The AlphaLISA assay development was performed in 96-well 12 area Alphaplate (reference 6052340, PerkinElmer, Waltham, MA) with a final reaction volume of 40 μL. Cross titration experiments of biotinylated MN (0 to 50 nM) against DIG-intasome (0 to 100 nM) were carried out to establish optimal assay concentrations for the binding assay. Ten μL of each protein were diluted in the binding buffer 50 mM BisTris-propane pH 7, 100 mM NaCl, 0.1 % (v/v) Tween-20, 0.1 % (w/v) BSA, 1 mM dithiothreitol. The plate was incubated at room temperature (RT) for 2 h in rotation. 10 μL of anti-DIG acceptor beads (PerkinElmer, reference AL113) were then added and after 1 h of incubation at RT with rotation, 10 μL of streptavidin donor acceptor beads (Perkin Elmer, reference 6760002) were mixed to the wells. This established a final concentration of 20 μg/mL for each bead. The plate was then incubated for 1 h at RT in the dark before the AlphaLISA signal was detected using an EnSpire Multimode Plate Reader (PerkinElmer). Negative control with binding buffer or only with one of the complexes were used to control the assay quality. Data were analyzed with GraphPad Prism 5.01 software. To evaluate tolerance for dimethyl sulfoxide (DMSO), the assay was performed as describe above with addition of 0.15 to 20 % (v/v) of DMSO during the binding step. For the competition experiment, increasing concentrations of untagged intasome were titrated out in the 1 nM DIG-intasome/biotinylated nucleosome interaction assay.

For the AlphaLISA screening assay with the NIH OncoSET library, the binding conditions were optimized for use in 384-well plate (Optiplate, reference 6007290, PerkinElmer) with the same concentrations for complexes and beads in a final reaction volume of 20.5 μL (2.5 % DMSO final concentration). Screening was first carried out with 25 μM of each compound pre-incubated for 1 h with 1 nM intasome/nucleosome complexes. Compounds were selected as inducing at least 50 % decrease of the AlphaLISA signal control condition and their inhibitory effect were further tested using dose-response curves.

For compounds validation, counter select assay was performed using short DNA fragment carrying a biotin and a DIG tag on each side. The AlphaLISA signal was monitoring using increasing concentrations of the double tagged DNA and the selected compounds were tested in optimized conditions at 10 μM. Any compound that causes decreased signal means that it interferes with AlphaLISA readout and therefore it is not relevant to this assay.

### MD simulation set-up and parametrisation

To evaluate the impact of the presence of doxorubicin on mononucleosome stability two models were prepared. One model of the nucleosome complex and another including the nucleosome complex plus 20 doxorubicin molecules.

The model for the nucleosome was prepared from the 5MLU structure (26), available in the Protein Databank (27), with a resolution of 2.80 Å. Protonation of all the amino acid residues was predicted using Propka version 3.0 at pH 7.0 (28). For the model with the doxorubicin molecules, packmol (29) was used to place and randomly distribute 20 doxorubicin molecules (1 for each 7 base pairs) around the volume defined by the two planes aligned with the enveloping DNA chains of the nucleosome.

Both systems were further prepared for molecular dynamics simulations using the AMBER18 software package and Xleap, using the ff14SB for the amino acid residues (30), the OL15 force field for DNA (31) and the General Amber Force Field (GAFF) for doxorubicin (32). Charges on the systems were neutralized through the addition of counter-ions and the system was placed in with TIP3P water boxes with a minimum distance of 12 Å between the nucleosome surface and the side of the box, using the LEAP module of AMBER.

Doxorubicin was parameterized with Antechamber using GAFF with RESP charges determined at HF/6-31G(d) using Gaussian16.

Both systems were submitted to four consecutive minimizations stages to remove clashes prior to the MD simulation. In these four stages the minimization procedure was applied to the following atoms of the system: 1-water molecules (2500 steps); 2-hydrogens atoms (2500 steps); 3-side chains of the amino acid residues and DNA (2500 steps); 4-full system (10.000 steps). The minimized systems were then subject to a molecular dynamics equilibration procedure, which was divided into two stages: In the first stage (50 ps), the systems were gradually heated to 310.15 K using a Langevin thermostat at constant volume (NVT ensemble); in the second stage (50 ps) the density of the systems was further equilibrated at 310.15 K.

Finally, molecular dynamic production runs were performed for 300 ns. These were performed with an NPT ensemble at constant temperature (310.15 K, Langevin thermostat) and pressure (1 bar, Berendsen barostat), with periodic boundary conditions, with an integration time of 2.0 fs using the SHAKE algorithm to constrain all covalent bonds involving hydrogen atoms. A 10 Å cutoff for nonbonded interactions was used during the entire molecular simulation procedure. Coordinates were saved at each 10 ps.

Final trajectories were analyzed in terms of backbone Root-mean-square deviation (RMSd), root-mean-square fluctuation (RMSF), and interactions formed.

### MM-GBSA

The molecular Mechanics / Generalized Born Surface Area method (33) was applied to estimate the histone-DNA binding free energies in the presence and in the absence of doxorubicin. From each MD trajectory, a total of 500 conformations taken from the last 200 ns of simulation were considered for each MM-GBSA calculation.

According to the MM-GBSA method the binding free energy can be decomposed as the sum of different energy terms, defined as:

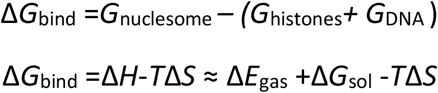

Because the structures of dimer and monomers or complex, protein, and ligand are extracted from the same trajectory, the internal energy change (Δ*E*_int_) is canceled.

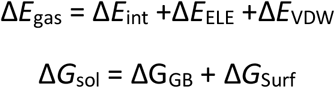

The gas-phase interaction energy (Δ*E*_gas_) between the components is written as the sum of electrostatic (Δ*E*_ELE_) and van der Waals (Δ*E*_VDW_) interaction energies. The solvation free energy (ΔG_sol_) is divided into the polar and non-polar energy terms. The polar solvation energy (ΔG_GB_) is calculated by using the Generalized-born (GB) model. In this case, the GB model proposed by Onufriev, Bashford and Case was considered (34). The non-polar contribution is calculated based on the solvent-accessible surface area (Δ*G*_Surf_), calculated in the present work with the LCPO method (35). The calculated binding free energy (Δ*G*_bind_) is hence written as the sum of the gas-phase interaction energy and solvation free energy.

### Infectivity assays

PBMC were isolated from blood samples using Ficoll-Hypaque gradient centrifugation. After separation, PBMC were pelleted by centrifugation. The cell culture medium consisted of RPMI 1640 supplemented with 20 % heat-inactivated fetal bovine serum, 5 % interleukin-2 (IL-2), and 50 μg gentamicin/ml. PBMC (2 × 10^6^) isolated from whole blood were incubated with various concentrations of RS-1 (0, 15, 30, 75, and 100 μM) for 24 h at 37°C. Next, PBMC were harvested (7 min, 400 × *g*) and resuspended in 500 μl culture medium. Ten microliters of HIV-1 subtype B virus (MOI = 0.1) was added to PBMC, and cells were incubated at 37°C for 3 h. Then, the medium was removed, and 10 ml of RPMI 1640 was added to wash the cells. The cells were harvested at low speed (400 × *g*), and the washed-cell pellet was resuspended in 2 ml of supplemented RPMI 1640. The cell suspension was added to wells of a 24-well tissue culture plate and incubated at 37°C. HIV-1 RNA from plasma samples was determined 24, 48, and 72 h postinfection. Replication in PBMC was quantified by HIV-1 RNA determination in cellular supernatant using Amplicor HIV Cobas TaqMan, version 2 (Roche, Basel, Switzerland), with a lower limit of detection of 20 copies/ml of plasma.

Viral DNA quantifications were performed as previously described (36). Cells were harvested at different time post infection by centrifugation of 2 × 10^6^- to 10 × 10^6^-cell aliquots, and cell pellets were kept frozen at −80°C until further analysis. Total DNA (including integrated HIV-1 DNA and episomal HIV-1 DNA) was extracted using the QIAmp blood DNA minikit (Qiagen, Courtaboeuf, France) according to the manufacturer’s protocol. Elution was performed in 50 μl of elution buffer. The total HIV-1 DNA was amplified by quantitative real-time PCR using the Light Cycler instrument (Roche Diagnostics, Meylan, France). Amplification was performed in a 20 μl reaction mixture containing 1× Light Cycler Fast Start DNA master hybridization probes (Roche Diagnostics), 3 mM MgCl_2,_ 500 nM forward primer LTR152 (5′-GCCTCAATAAAGCTTGCCTTGA-3′, and 500 nM reverse primer LTR131 (5′-GGCGCCACTGCTAGAGATTTT-3′), located in an LTR region with highly conserved fluorogenic hybridization probe LTR1 (50 nM; 5′-6-carboxyfluorescein [FAM]-AAGTAGTGTGTGCCCGTCTGTT[AG]T[GT]TGACT-3′-6-carboxytetramethylrhodamine [TAMRA]). After an initial denaturation step (95°C for 10 min), total HIV-1 DNA was amplified for 45 cycles (95°C for 10 s, 60°C for 30 s), followed by 1 cycle at 40°C for 60 s. The copy number of total HIV-1 DNA was determined using the 8E5 cell line. The 8E5/LAV cell line, used for a standard curve, was derived from a CEM cellular clone containing a single, integrated, defective (in the *pol* open reading frame), constitutively expressed viral copy. 8E5 DNA (5 to 5 × 10^4^ copies) was amplified. Results were expressed as the copy number of total HIV-1 DNA per 10^6^ cells.

The 2-LTR DNA circles were amplified with primers HIV-F and HIV-R1, spanning the LTR-LTR junction, as described elsewhere (37). Briefly, amplification was performed in a 20-μl reaction mixture containing 1× Light Cycler Fast Start DNA master hybridization probes (Roche Diagnostics), 4 mM MgCl_2_, 300 nM forward and reverse primers spanning the LTR-LTR junction, and 200 nM each fluorogenic hybridization probe. Copy number of 2-LTR circles was determined in reference to a standard curve prepared by amplification of quantities ranging from 10 to 1 × 10^6^ copies of a plasmid comprising the HIV_LAI_ 2-LTR junction (37) by using Light Cycler quantification software, version 4.1 (Roche Diagnostics). Results are expressed as copy number of 2-LTR circles per 1× 10^6^ cells.

Integrated DNA was first amplified by Alu PCR performed in a 50-μl reaction mixture containing 200 ng total DNA, 1× HF Phusion mix, 200 nM deoxynucleoside triphosphates (dNTP), 500 nM primer PBS (5′-TTTCAAGTCCCTGTTCGGGCGCCA-3′), located in the PBS sequence of the viral genome, and 500 nM primer Alu-164 (5′-TCCCAGCTACTGGGGAGGCTGAGG-3′), located in the Alu sequence of the cellular genome. After an initial denaturation step (98°C for 30 s), the heterogeneously sized population of integrated DNA was amplified for 35 cycles (98°C for 10 s, 60°C for 20 s, 72°C for 2 min 30 s). A second nested PCR was then performed in a 50-μl reaction mixture containing 5 μl of Alu PCR product, 1× HF Phusion mix, 500 nM dNTP, 500 nM primer NI-1 (5′-CACACACAAGGCTACTTCCCT-3′), and 500 nM primer NI-2 (5′-GCCACTCCCCAGTCCCGCCC-3′); primer sequences match sequences localized in the viral genome. After an initial denaturation step (94°C for 12 min), the expected 351-bp fragment of integrated DNA was amplified for 42 cycles (94°C for 1 min, 60°C for 1 min, 72°C for 1 min). Results were analyzed on 1.2% agarose SYBR safe stain gel. A first-round PCR control was ran in the absence of polymerase in order to quantify any unspecific amplification during the second round. The copy number of integrated HIV-1 DNA was determined in reference to a standard curve generated by concomitant two-stage PCR amplification of a serial dilution of the standard HeLa R7 Neo cell DNA mixed with uninfected-cell DNA to yield 50,000 cell equivalents. Cell equivalents were calculated according to amplification of the β-globin gene (two copies per diploid cell) with commercially available materials (Control Kit DNA; Roche Diagnostics). 2-LTR circles, total and integrated HIV-1 DNA levels were determined as copy numbers per 10^6^cells. Two-LTR circles and integrated cDNA were also expressed as a percentage of the total viral DNA.

## DATA AVAILABILITY

All data are available upon request.

## AKNOWLEDGEMENTS

English editing has been performed by Nature Rearch Editing Service.

## SUPPLEMENTARY DATA

**Sup data 1.**
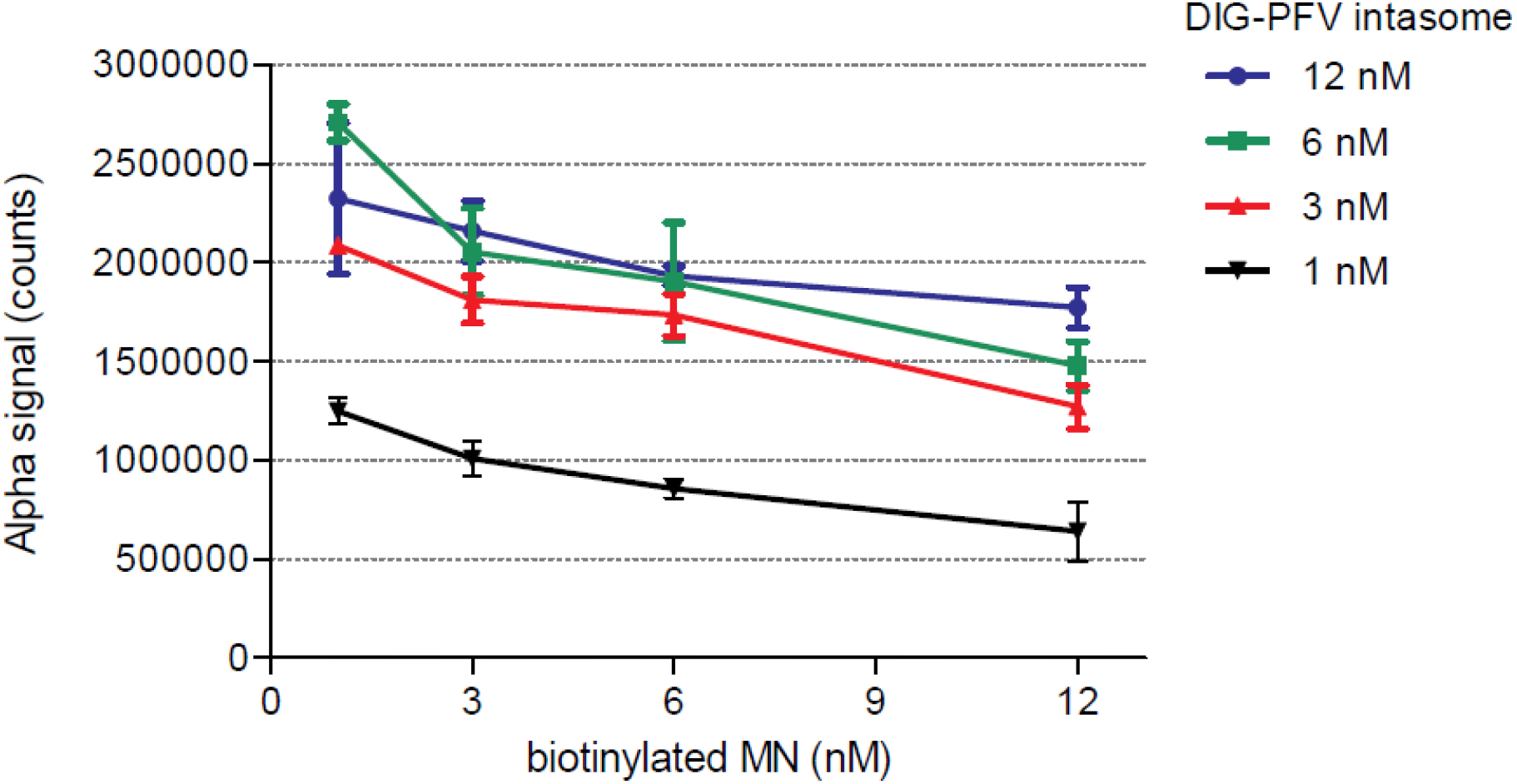
AlphaLisa cross titration assay. The Alphascreen interaction signal (AU) was monitored using increasing concentration of each partners and data are reported as mean of three independent experiments ±SD.

**Sup data 2.**
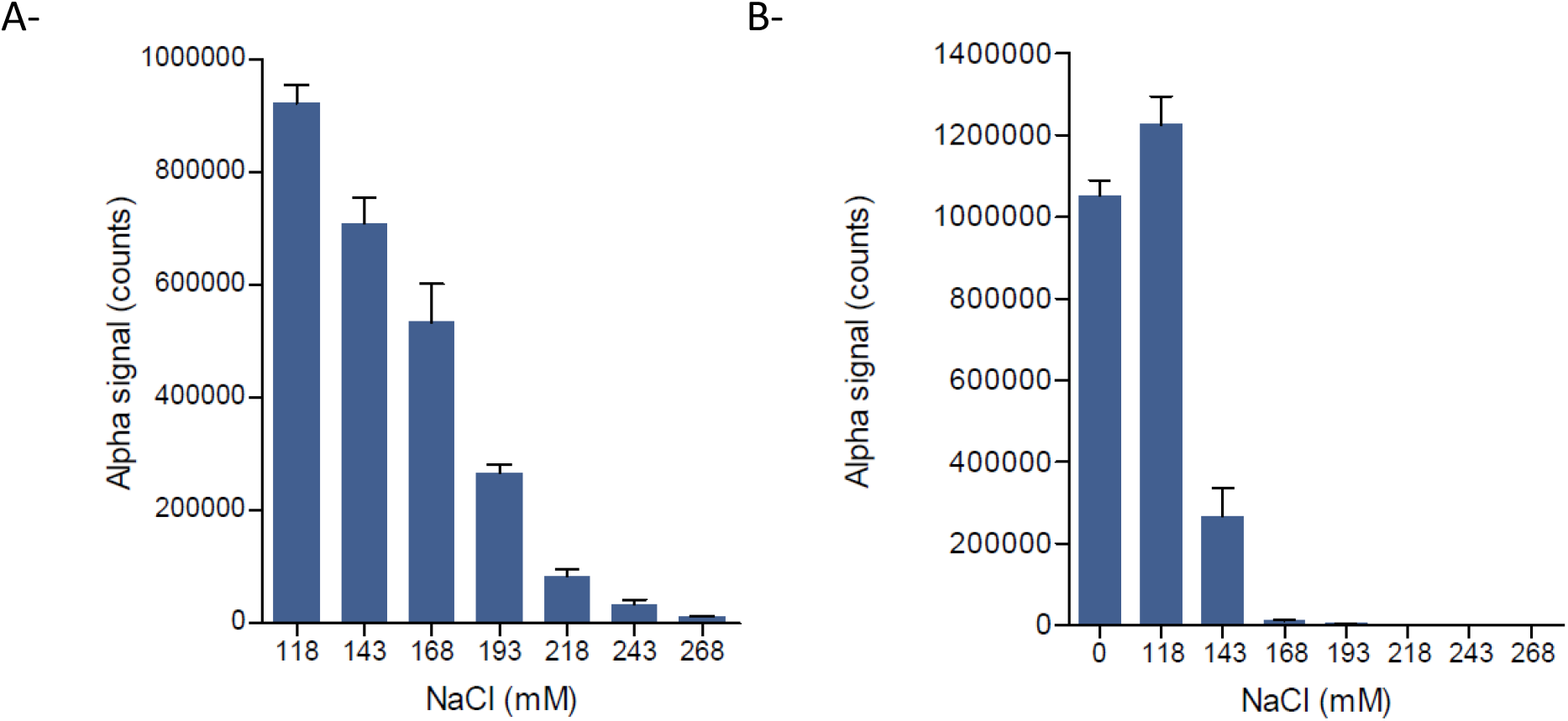
Effect of salt concentration on PFV intasome binding to the nucleosome in AlphaLisa assay. The Alphascreen interaction signal (AU) was monitored using increasing concentration of NaCl using 3nM of intasome and 3nM of either nucleosome (**A**) or naked DNA (**B**). Data are reported as mean of three independent experiments ±SD.

**Sup data 3.**
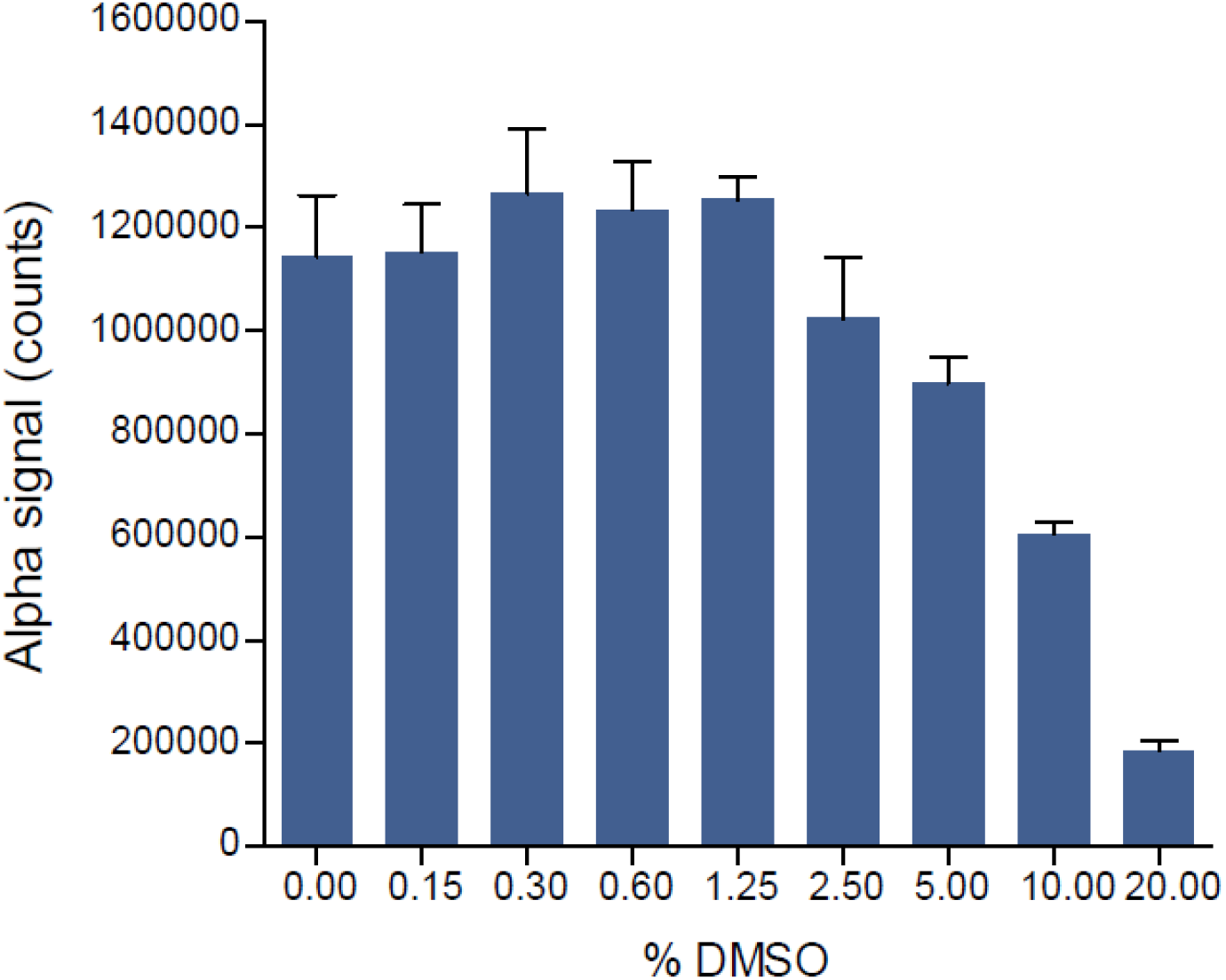
Effect of DMSO concentration on PFV intasome binding to the nucleosome in AlphaLisa assay. The Alphascreen interaction signal (AU) was monitored using increasing concentration of DMSO using 3nM of each partner. Data are reported as mean of three independent experiments ±SD.

**Sup data 4.**
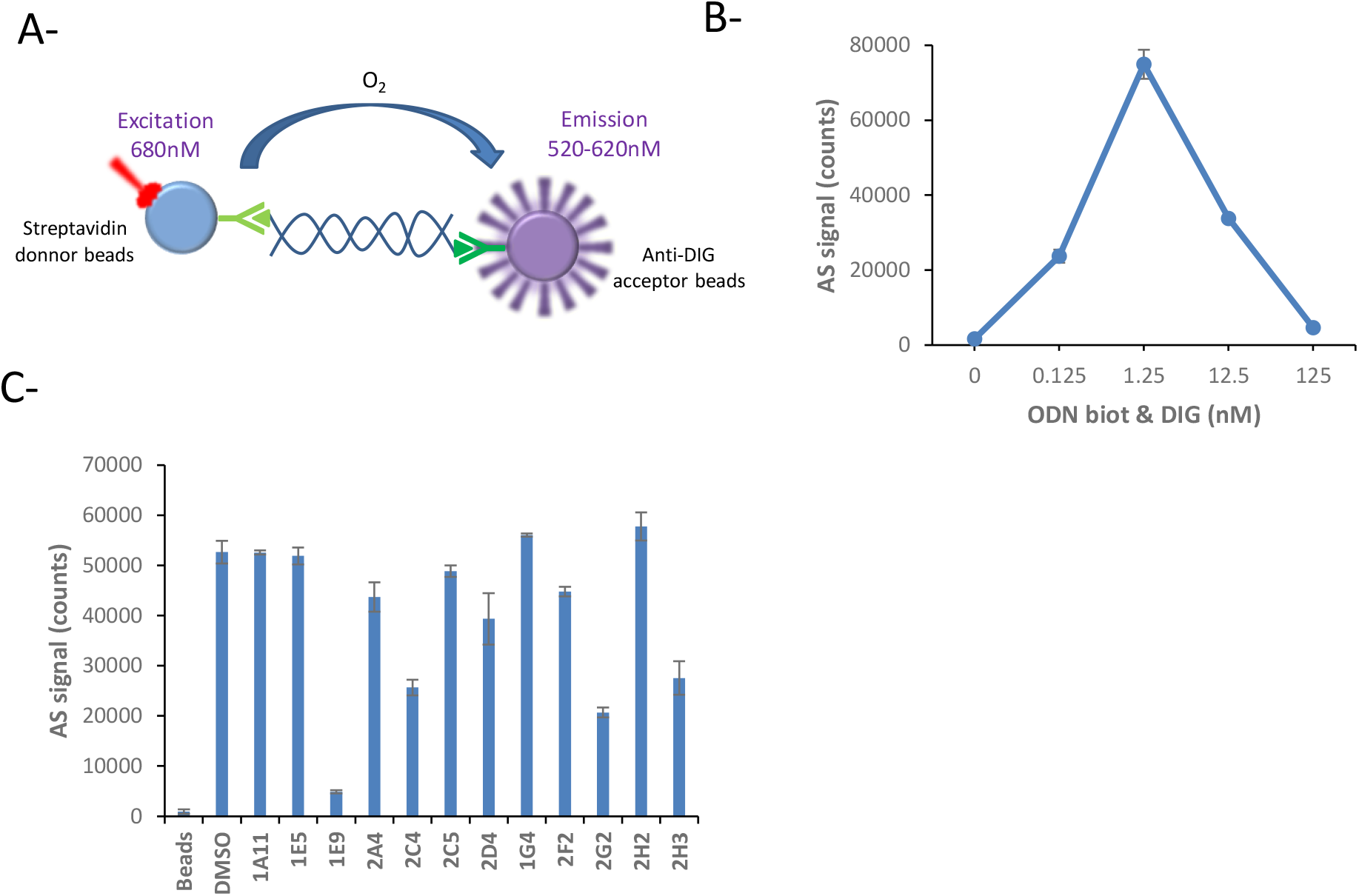
Counter select assay. The counter select assay was performed using short DNA fragment carrying a biotin and a DIG tag on each side (**A**). The Alphascreen interaction signal (AU) was monitored using increasing concentration the double tagged DNA (**B**). Each selected compound was then tested in optimized conditions at 10μM. Data are reported as mean of three independent experiments ±SD.

**Sup data 5.**
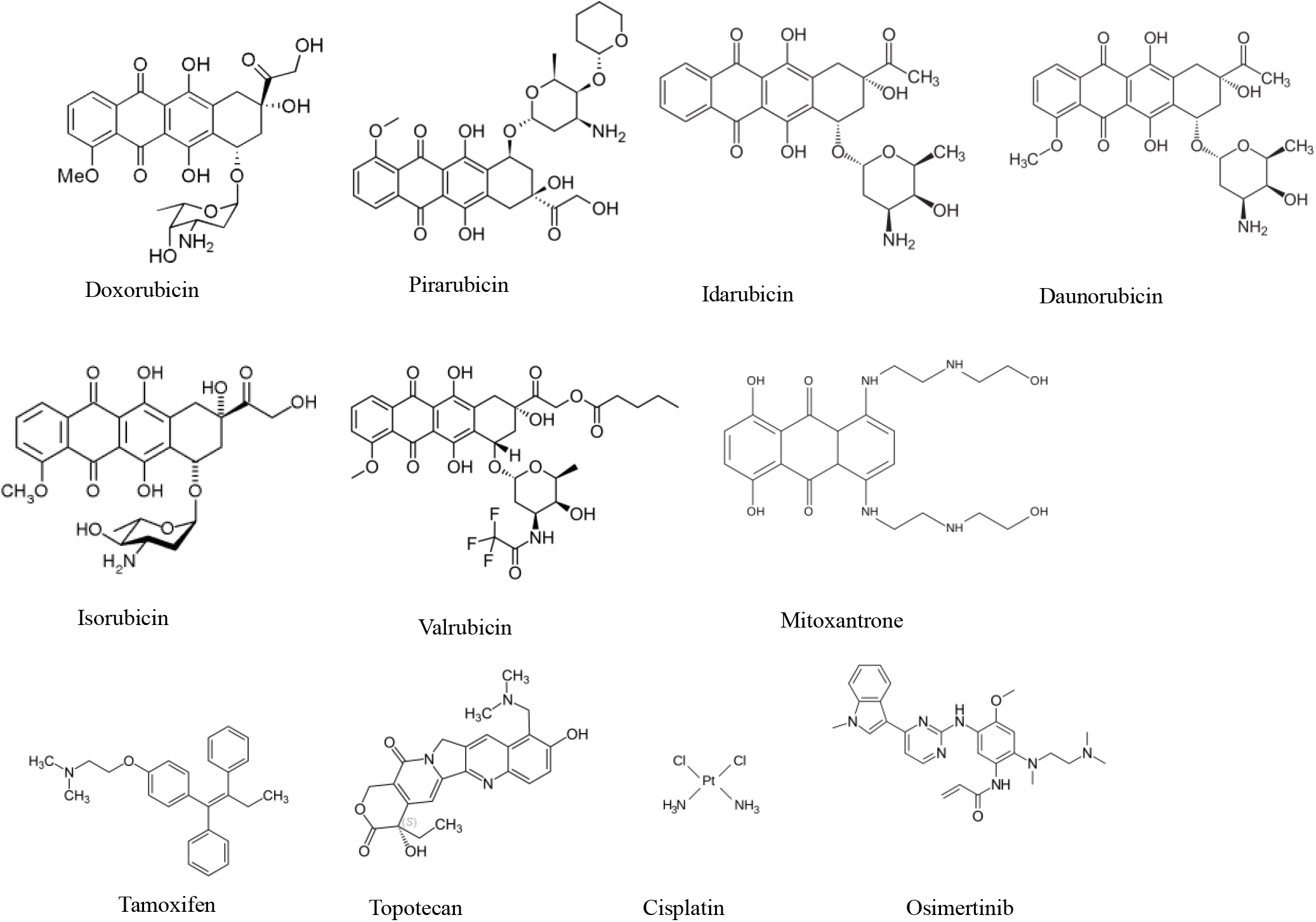
Chemical structures of the ONCOSET drugs used in this work.

**Sup data 6.**
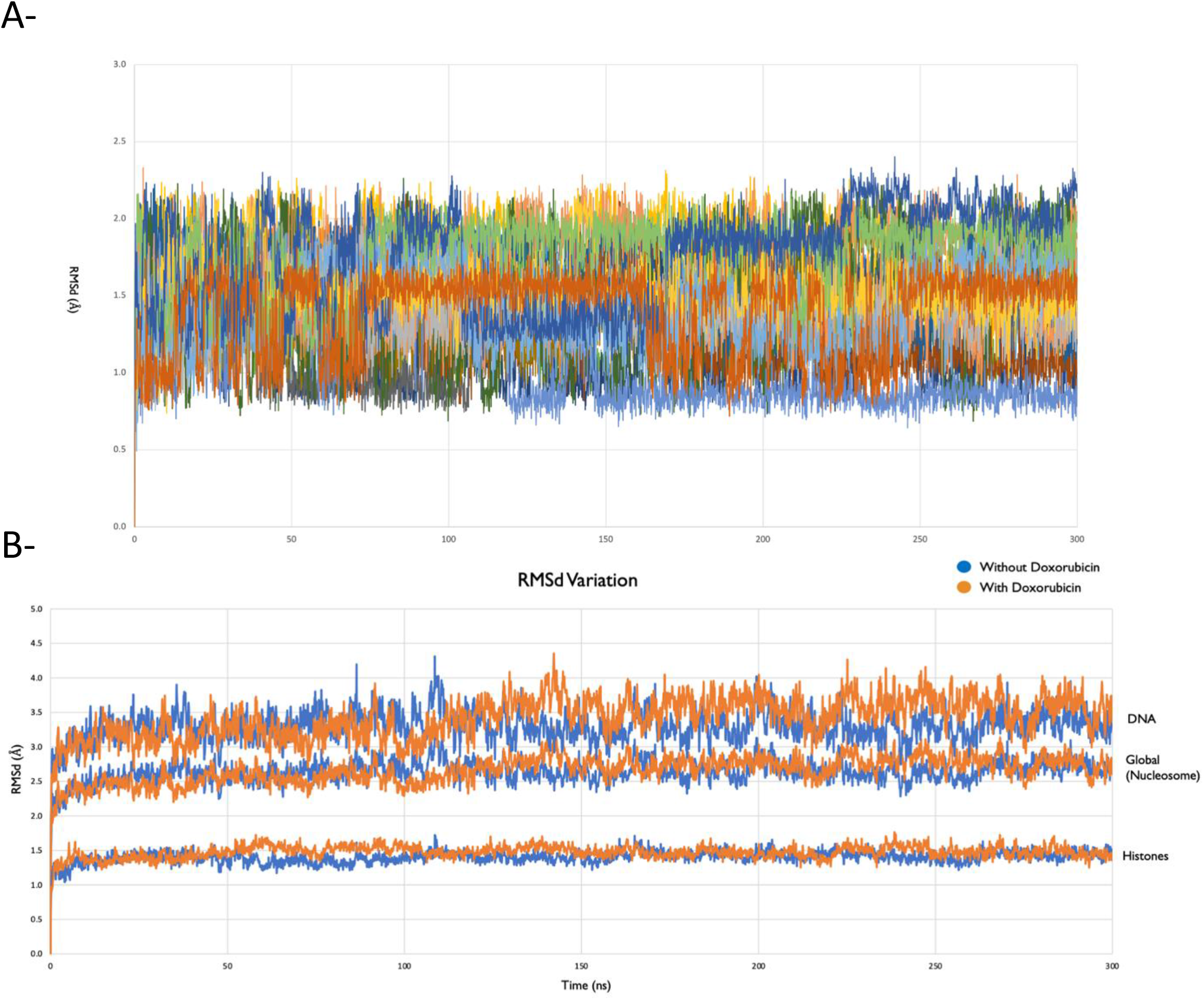
A-Representation of the RMSd for each of the doxorubicin molecules considered in the nucleosome simulations. SI6 B-RMSd variation of the backbone atoms of the nucleosome along the MD simulations.

**Sup data 7.**
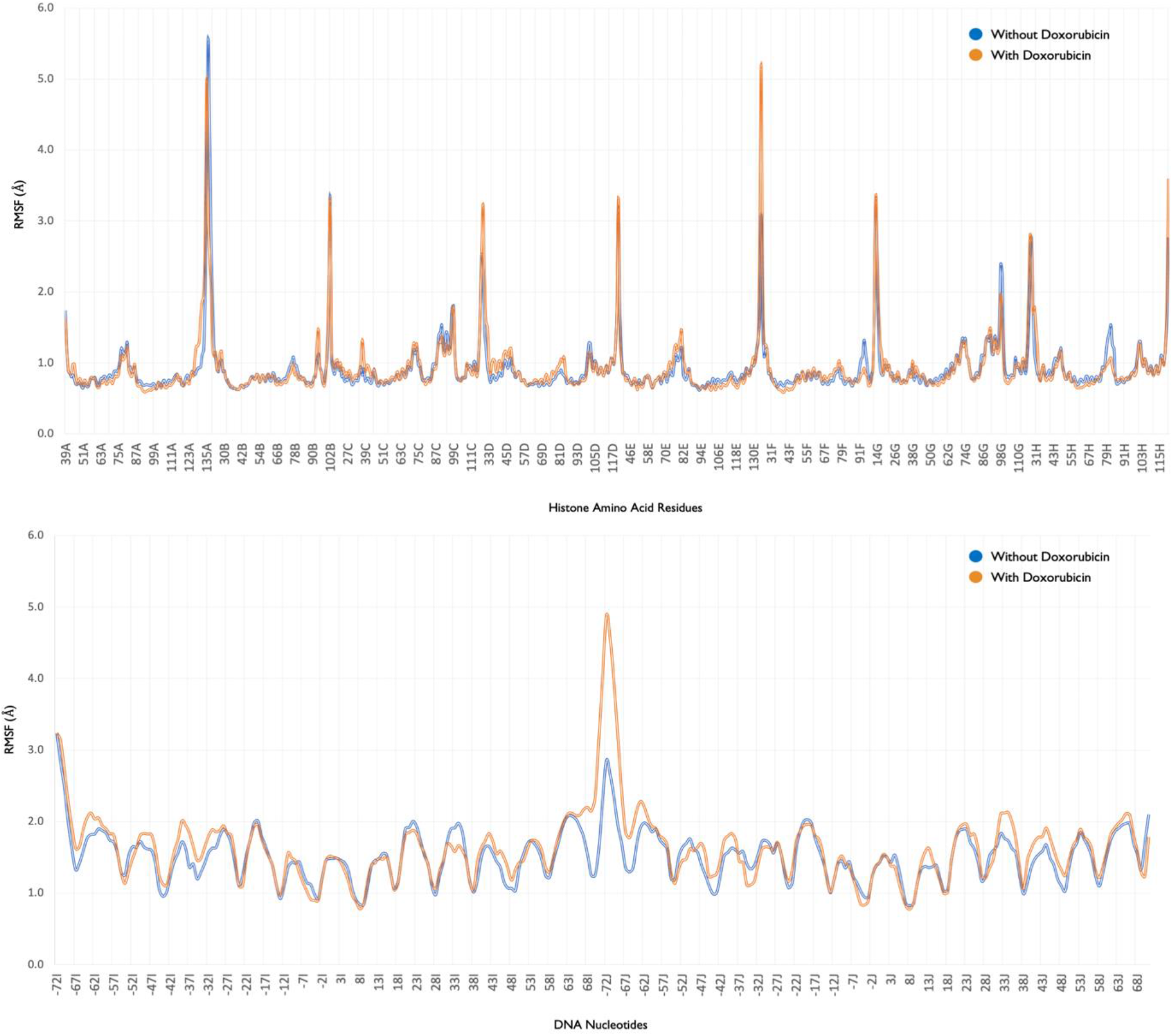
RMSF variation of the different residues and nucleotides of the nucleosome in the simulations done in the presence and absence of doxorubicin.

**Sup data 8.**
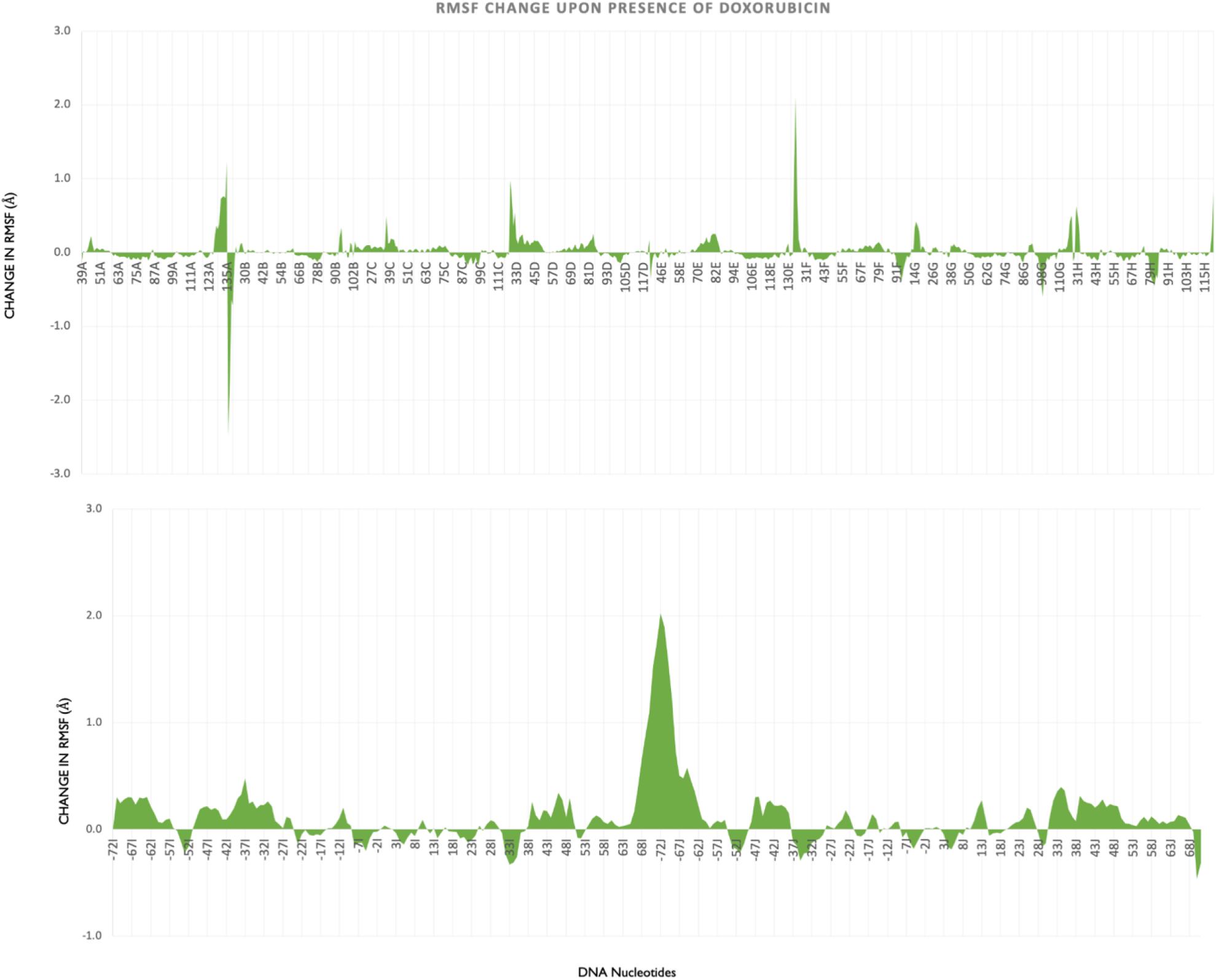
Change in RMSF upon addition of doxorubicin for the histone amino acid residues and DNA nucleotides.

**Sup data 9.**
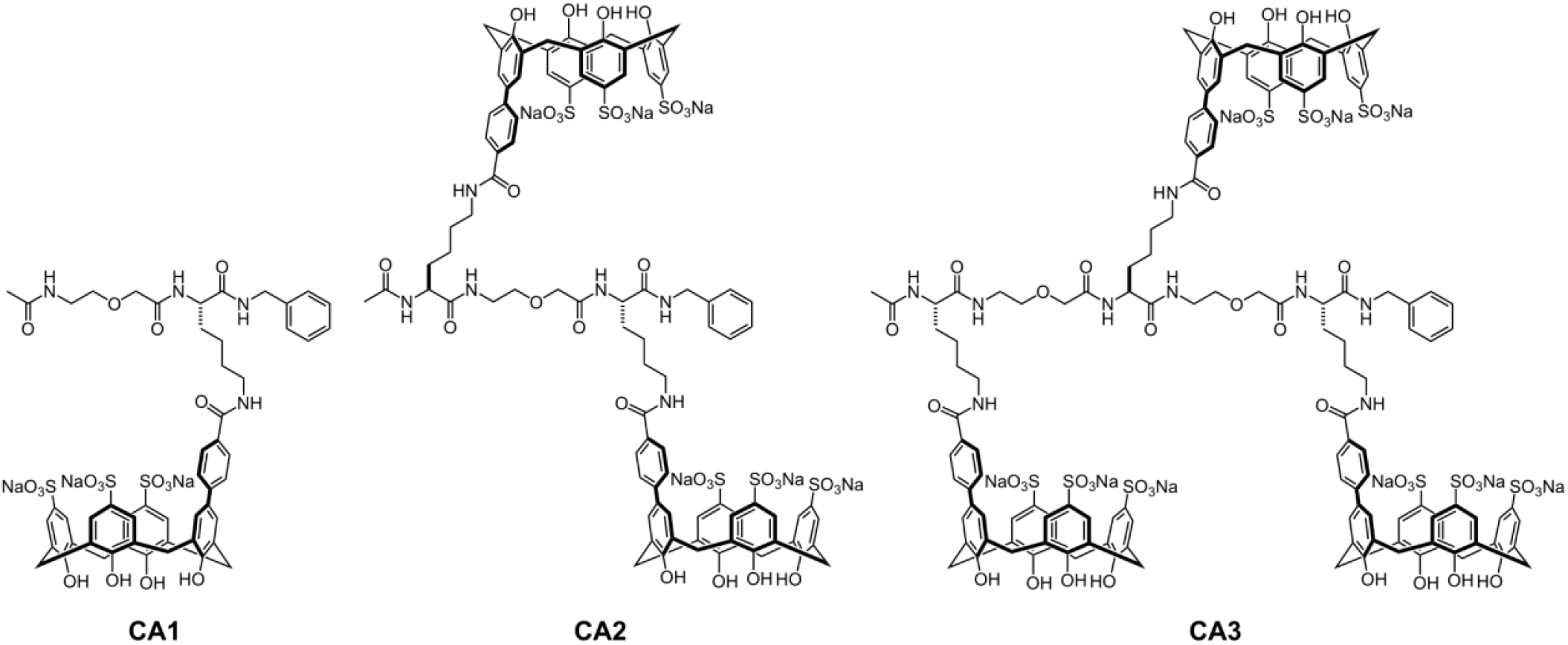
Chemical structure of calixarenes molecules CA1, CA2 and CA3.

**Sup data 10.**
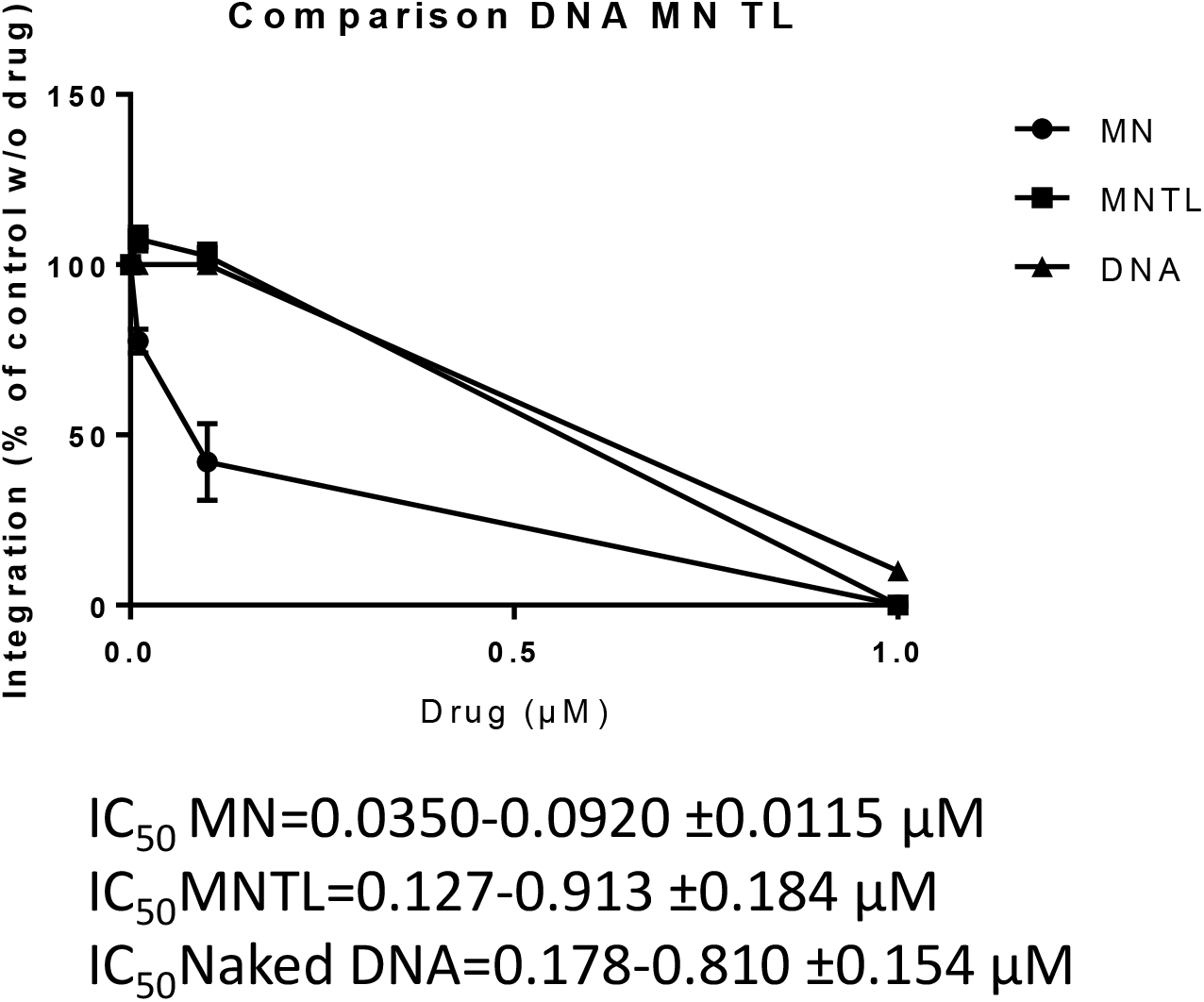
Effect of CA3 molecules on *in vitro* concerted integration catalyzed by HIV-1 intasomes on mononucleosome (MN), tailless nucleosome (MNTL) or naked 601 DNA. CA3 (**C, D**) has been added to a typical *in vitro* concerted integration performed with HIV-1 IN, LEDGF/p75 cofactor, radiolabeled viral U5 end fragment and MN, MNTL or naked DNA. The integration products were monitored on 6-12 % gradient polyacrylamide gel and quantified. Data are reported as mean of 3-4 independent experiments ±SD and IC_50_ were reported in the figure.

**Sup data 11.**
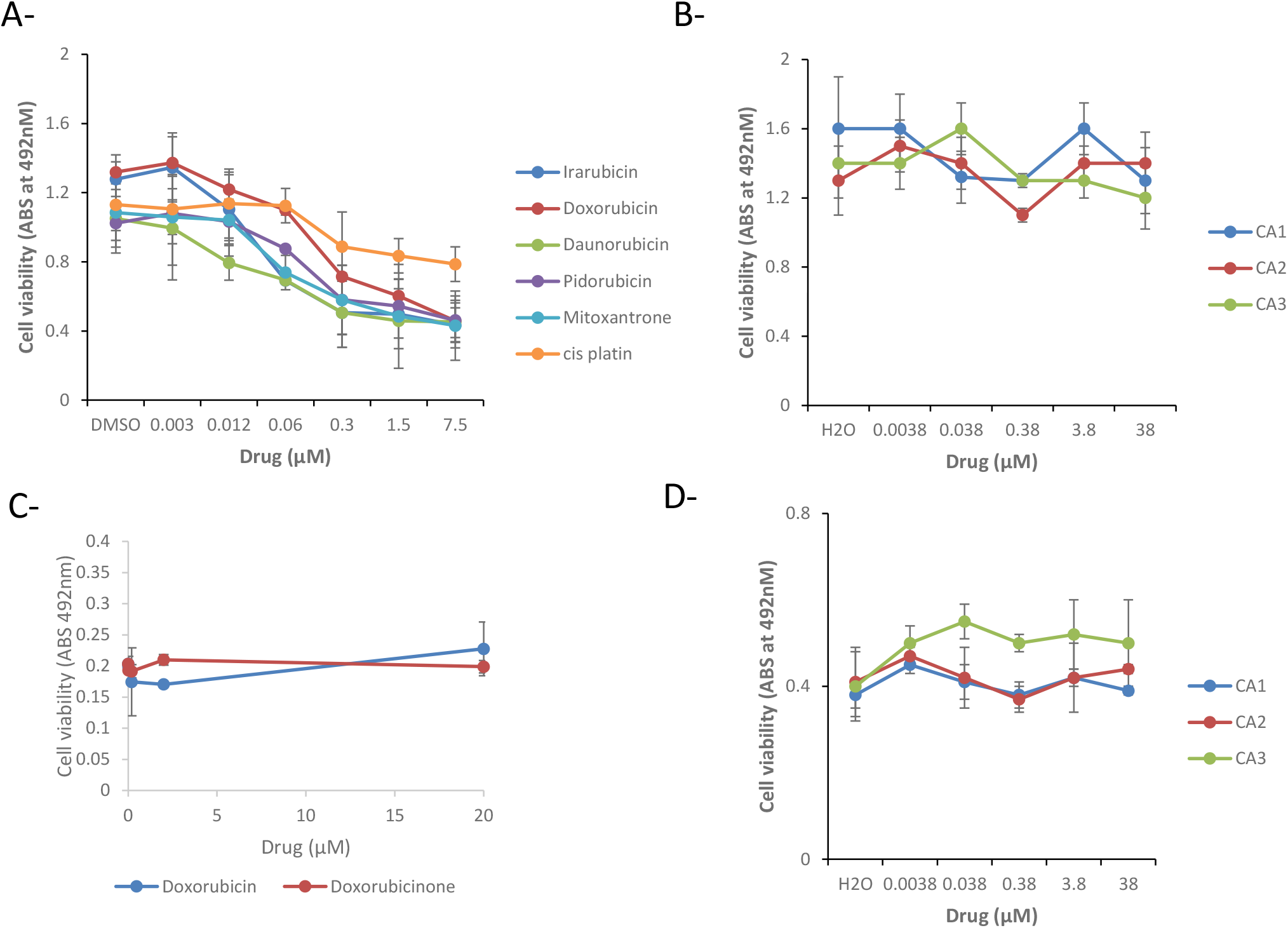
Cell toxicity of the drugs were tested in HEK293T (A-B) and PBMC cells (C-D). Cells were treated 24h and viability was measured by MTT assay as indicated in Material and Methods section. Absorbance was measured at 492 nm and data are reported as mean of 3 independent experiments ±SD.

